# Corticothalamic gating of population auditory thalamocortical transmission in mouse

**DOI:** 10.1101/625988

**Authors:** Baher A. Ibrahim, Caitlin Murphy, Guido Muscioni, Aynaz Taheri, Georgiy Yudintsev, Robert V. Kenyon, Tanya Berger-Wolf, Matthew I. Banks, Daniel A. Llano

## Abstract

Since the discovery of the receptive field, scientists have tracked receptive field structure to gain insights about mechanisms of sensory processing. At the level of the thalamus and cortex, this linear filter approach has been challenged by findings that populations of cortical neurons respond in a stereotyped fashion to sensory stimuli. Here, we elucidate a possible mechanism by which gating of cortical representations occurs. All-or-none population responses (here called “ON” and “OFF” responses) were observed in vivo and in vitro in the mouse auditory cortex at near-threshold acoustic or electrical stimulation. ON-responses were associated with previously-described UP states in the auditory cortex. OFF-responses in the cortex were only eliminated by blocking GABAergic inhibition in the thalamus. Opto- and chemogenetic silencing of NTSR-positive corticothalamic layer 6 (CTL6) neurons as well as the pharmacological blocking of the thalamic reticular nucleus (TRN) retrieved the missing cortical responses, suggesting that the corticothalamic feedback inhibition via TRN controls the gating of thalamocortical activity. Moreover, the oscillation of the pre-stimulus activity of corticothalamic cells predicted the cortical ON vs. OFF responses, suggesting that underlying cortical oscillation controls thalamocortical gating. These data suggest that the thalamus may recruit cortical ensembles rather than linearly encoding ascending stimuli and that corticothalamic projections play a key role in selecting cortical ensembles for activation.

## Introduction

Our interactions with the world depend on how the sensory information is transmitted, integrated, and processed in the nervous system. Most models of perception propose that activation of the cerebral cortex is critical for conscious experience of sensory stimuli. For most of our senses, the thalamus is a critical brain structure to allow information to reach the cortex. One view of thalamic function proposed that thalamus governs the sensory representation in the cortex by linearly transmitting the sensory information from lower sensory structures to the cortex. This view emerged after the descriptions of receptive field transformations in the visual system by Hubel and Wiesel (*1*). According to this idea, cortical activity patterns during sensory perception should be predictable based on activity patterns in the thalamus and patterns of synaptic convergence of thalamocortical neurons onto cortical neurons.

However, this view does not comport with findings that population activity in sensory cortices is often stereotyped and recapitulates patterns of cortical spontaneous activity (*2–5*). In addition, a linear filter model cannot explain the presence of formed complex hallucinations, which are associated with elevated activity in the primary sensory cortices (*6–8*). As such, a hypothesis has emerged that sensory representations are developed by early exposure to sensory stimuli and stored in the cortex in intracortical networks, and that the thalamus activates these pre-wired sensory representations upon sensory stimulation (*2, 9*). Some observations could support this hypothesis. For example, ongoing cortical activity which was reported *in-vivo* and *in-vitro* (*4, 10*)was found to be highly determined by the internal cortical connectivity (*4, 11–13*), totally independent on the thalamocortical afferents (*2, 14, 15*), and is the main platform for the generation of the internal percepts in memory and REM sleep (*16, 17*). In addition, despite substantial differences in the form and organization in the initial processing stages of different modalities of sensation, at the level of the thalamus and cortex, neural circuits across modalities are strikingly similar. These data suggest that there is a common function of thalamocortical circuits that is not tied to specific modalities of perception. Given that connected neuronal ensembles could be the main functional unit for behavior and cognition (*18, 19*), the ability of thalamocortical afferents to activate the same cortical circuits suggests that a major control point for the activation of cortical ensembles lies in the thalamus. Here, we examine the mechanisms of gating all-or-none population responses in the auditory cortex (AC) and show that ongoing oscillatory activity in layer 6 corticothalamic projections gates the activation of cortical ensembles. The gating of cortical activity occurs via corticothalamic projection to the thalamic reticular nucleus (TRN), which is a long-enigmatic structure that partially surrounds and sends GABAergic projections to thalamocortical neurons. These findings suggest that at least one mode of thalamocortical function is to select particular groups of cortical neurons for activation based on feedback from corticothalamic neurons.

## Material and Methods

### Animals

C57BL/6J (Jackson Laboratory, stock # 000664), C57BL/6J-Tg (Thy1-GCaMP6s) GP4.3Dkim/J a.k.a. GCaMP6s mice (Jackson Laboratory, stock # 024275), BALB/cJ (Jackson Laboratory, stock # 000651), Gad2-IRES-Cre (Jackson Laboratory, stock # 010802), NTSR1-Cre (a generous gift from Dr. Gordon Shepherd from Northwestern University), and RBP4-Cre (received from cryopreserved stock from the Mutant Mouse Resource and Research Center (MMRRC, stock number 031125-UCD)) mice of both sexes were used. All applicable guidelines for the care and use of animals were followed. All surgical procedures were approved by the Institutional Animal Care and Use Committee (IACUC) at University of Illinois Urbana-Champaign. Animals were housed in animal care facilities approved by the American Association for Accreditation of Laboratory Animal Care (AAALAC).

### *In-vivo* imaging

The detailed procedures were described before [(*20*)]. In brief, GCaMP6s mice were used for transcranial *in-vivo* imaging of evoked calcium signals from the left auditory cortex (AC). For each experiment, the mouse was anesthetized with a mixture of ketamine and xylazine (100 mg/kg and 3 mg/kg respectively) delivered intraperitoneally. The animal’s body temperature was maintained within the range of 35.5 and 37 °C using a DC temperature controller (FHC, ME, USA). A mid-sagittal and mid-lateral incisions were made to expose the dorsal and lateral aspects of the skull along with the temporalis muscle. The temporalis muscle was separated from the skull to expose the ventral parts of the underlying AC. The site was cleaned with sterile saline, and the surface of the skull was thinned by a specific drill. A small amount of dental cement (3M ESPE KETAC) was mixed to a medium level of viscosity and added to the head of the bolt just enough to cover it. The bolt was bonded to the top of the skull, and the dental cement was allowed to set.

An Imager 3001 Integrated data acquisition and analysis system (Optical Imaging Ltd., Israel) was used to image the cortical responses to sound in mice. A macroscope consisting of 85 mm f/1.4 and 50 mm f/1.2 Nikon lenses were mounted to an Adimec 1000m high-end CCD camera (7.4 x 7.4-pixel size, 1004 × 1004 resolution), and centered above the left AC, focused approximately 0.5 mm below the surface of the exposed skull. Acoustic stimuli were generated using a TDT system 3 with an RP 2.1 Enhanced Real-Time Processor and delivered via an ES1 free field electrostatic speaker (Tucker-Davis Technologies, FL, USA), located approximately 8 cm away from the contralateral ear. All imaging experiments were conducted in a sound-proof chamber at 10 frames per second. 500 ms pure tones of 5 kHz, 37 dB SPL, 100% amplitude modulated at 20 Hz were played every 10 seconds. ΔF/F of evoked calcium signals following the sound presentation was obtained.

### Virus injection

To modulate specific cell types of animal’s brain, Cre-technology was used to provide an expression of opto- or chemo-genetic probes in those specific cells after 11 days of viral injection at P4. The detailed procedures were described before [(*21*), in press]. For all neonates, cryoanesthesia was induced after five to ten minutes. A toe pinch was done to confirm that the mice were fully anesthetized. A small animal stereotaxic instrument (David Kopf Instruments, Tujunga, CA) was used with a universal syringe holder (David Kopf Instruments, Tujunga, CA) and standard ear bars with rubber tips (Stoelting, Wood Dale, IL). The adaptor stage was cooled by adding ethanol and dry ice to the well. A temperature label (RLC-60-26/56, Omega, Norwalk, CT) was attached to the adaptor to provide the temperature of the stage during cooling. The temperature was kept above 2°C to prevent hypothermia or cold-induced skin damage of the neonatal mice and below 8°C to sustain cryoanesthesia. Glass micropipettes (3.5-inches, World Precision Instruments, Sarasota, FL) were pulled using a micropipette puller (P-97, Sutter Instruments, Novato, CA) and broken back to a tip diameter between 35-50μm. The micropipette was filled with mineral oil (Thermo Fisher Scientific Inc., Waltham, MA) and attached to a pressure injector (Nanoliter 2010, World Precision Instruments, Sarasota, FL) connected to a pump controller (Micro4 Controller, World Precision Instruments, Sarasota, FL). To target corticothalamic L6, layer 5 (L5), the AC of NTSR1-Cre (*22–25*) or RPB4-Cre (*26, 27*) neonates was injected with “eNpHR3.0” AAV1 (AAV-EF1a-DIO-eNpHR3.0-YFP) (Halorhodopsin-AAV) constructs from UNC Vector Core (Chapel Hill, NC) or Gi-coupled hM4Di DREADDs AAV8 (AAV8-DIO-hSyn-hM4Di-mCherry) (DREADDs-AAV) construct from Addgene (Cambridge, MA). The micropipette carrying the viral particles was first located above the AC at the left hemisphere at 1.5 mm anterior to lambda and just at the edge of the skull’s flat horizon. The tip was lowered to 1.2 mm from the brain surface, then it was pulled back to 1.0 mm for the first injection where 200 nL of Halorhodopsin-AAV or DREADDs-AAV was injected at 200 nL/min. After the injection was finished, the micropipette was left in the brain for 1 minute before removing to allow the injectate to settle into the brain. Following the first injection, the tip was pulled back stepwise in 0.1 mm increments, and 200 nL of the injectate was injected at every step until the tip reached 0.3mm from the surface. In total, 1600 nL of AAV was injected into the AC. The incision was sutured using 5/0 thread size, nylon sutures (CP Medical, Norcross, GA). To target the GABAergic cells of the inferior colliculus (IC), the IC of GAD2-Cre (*28–30*) neonatal mice was injected with Ha-AAV following the same procedures showing above, but the micropipette loaded by Ha-AAV was located over the IC at the left hemisphere at 2.0 mm posterior to lambda and 1.0 mm laterally from the midline. The neonates were transferred back onto a warming pad to recover. After 5-7 minutes, their skin color was returned to normal and they started moving. After recovery, all neonates were returned to their nest with the parents.

### Brain slicing

For all *in-vitro* experiments, 15-18 days old mice were initially anesthetized with ketamine (100 mg/ kg) and xylazine (3 mg/kg) intraperitoneally and transcardially perfused with chilled (4 °C) sucrose-based slicing solution containing the following (in mM): 234 sucrose, 11 glucose, 26 NaHCO_3_, 2.5 KCl, 1.25 NaH_2_PO_4_, 10 MgCl_2_, 0.5 CaCl_2_. After the brain was taken out, it was cut to obtain auditory colliculo-thalamocortical brain slice (aCTC) as shown (Figure S1) and as described before (*31–33*). 600 μm thick horizontal brain slices were obtained to retain the connectivity between inferior colliculus (IC), medial geniculate body (MGB), thalamic reticular nucleus (TRN) and AC. All slices were incubated for 30 min in 33 °C in a solution composed of (in mM: 26 NaHCO_3_, 2.5 KCl, 10 glucose, 126 NaCl, 1.25 NaH_2_PO_4_, 3 MgCl_2_, and 1 CaCl_2_). After incubation, all slices were transferred to a perfusion chamber coupled to an upright Olympus BX51 microscope, perfused with artificial cerebrospinal fluid (ACSF) containing (in mM) 26 NaHCO_3_, 2.5 KCl, 10 glucose, 126 NaCl, 1.25 NaH_2_PO_4_, 2 MgCl_2_, and 2 CaCl_2_. Another set of experiments was done at a different lab [Dr. Matthew I. Banks (Madison, WI)] to exclude any experimental factors related to our lab environment, chemicals, or anesthesia. As reported by the lab (*34*), following full anesthesia by isoflurane, C57BL/6J mouse was immediately decapitated without cardiac perfusion, the animal’s brain was extracted and immersed in cutting artificial CSF [cACSF; composed of (in mM) 111 NaCl, 35 NaHCO_3_, 20 HEPES, 1.8 KCl, 1.05 CaCl_2_, 2.8 MgSO_4_, 1.2 KH_2_PO_4_, and 10 glucose] at 0–4°C. Slices were maintained in cACSF at 24°C for >1 h before transfer to the recording chamber, which was perfused at 3–6 ml/min with ACSF [composed of (in mM) 111 NaCl, 35 NaHCO_3_, 20 HEPES, 1.8 KCl, 2.1 CaCl_2_, 1.4 MgSO_4_, 1.2 KH2PO_4_, and 10 glucose]. All the solutions were bubbled with 95% oxygen/5% carbon dioxide.

### Electrical stimulation

All the electrical stimulation protocols in the IC evoked the same cortical response modes. Following the electrical stimulation of IC, the first step was always to get a neuronal activation in all brain structures involved in the auditory circuit to make sure that aCTC slice retains the synaptic connection between IC, MGB, TRN, and AC. For imaging the cortical activation, one second electrical train pulses of (250uA, 40Hz, 1ms pulse width) to IC was the main stimulating protocol as described before (*31*). However, one second long of electrical stimulation was not suitable for electrophysiology experiments to record any post-stimulus signals that could be buried by the stimulus artifact. Other IC stimulating protocols were used. For whole cell and LFP recording, the stimulation of IC by (3 ms long train pulses, 300-500 uA, 1000 Hz, 1ms pulse width) or (100 ms long train pulses, 250uA, 40 Hz, 1ms pulse width) was used. The electrical stimulation was done by a concentric bipolar electrode (Cat#30201, FHC) every 10-20s. The parameters of the electrical pulses were adjusted by a B&K precision wave generator (model # 4063) and World Precision Instruments stimulation isolator (A-360).

### Calcium (Ca) and flavoprotein (FA) Imaging

For Ca imaging, GCaMP6s mouse or loading CAL-520, AM (Cal-520, AM (Abacam, ab171868) calcium dye was used. For CAL-520, AM calcium dye loading, the brain slicing protocol was followed, but the aCTC slices were incubated in (48 ul of DMSO dye solution, 2 ul of Pluronic F-127 (Cat# P6866, Invitrogen), and 2.5 ml of the incubating solution) at 35-36 °C for 25-28 minutes according to (R). The slices then were incubated in the normal incubating solution (shown above) for 30 minutes to wash the extra extracellular dye. Imaging was done under ACSF perfusion as described before. Depending on the experiment, the evoked Ca or FA signals following IC stimulation were tracked using a stable DC fluorescence illuminator (Prior Lumen 200) and a U-M49002Xl E-GFP Olympus filter cube set [excitation: 470–490 nm, dichroic 505 nm, emission 515 nm long pass, 100 ms exposure time for FAD and 5 ms for calcium signals]. All data were collected using Retiga EXi camera of a frame rate as 4 Hz for FAD and 10 Hz for Ca imaging. The peak of the signals was detected by placing ROI on the brain regions (IC, MGB, or AC) or the individual cortical or thalamic cells. ΔF/F of FA responses from IC, MGB, AC was obtained for further analysis.

### Pharmacological intervention

To disinhibit the inhibitory inputs globally, GABAα-R antagonist (*35*), SR-95531 (gabazine, Cat# 1262, Tocris) was added to the perfused ACSF solution (200 nM). To specifically disinhibit the inhibitory inputs in either MGB or AC, a continuous flow of gabazine (200 nM) was injected through a glass pipette (Broken tip, 35 uM) which was connected to a picospritzer (TooheyCompany, New Jersey, USA). The pipette was filled by a solution composed of (1ml ACSF+10ul Alexa Fluor 594 hydrazide, sodium salt dye, Cat#A10438, Invitrogen). The dye was used to visualize the flow of the solution and to make sure it is only going to the site of injection. The injection was done under 10 psi pressure for 5 minutes and continuously during imaging. As reported before, to block TRN activity (*36, 37*), AMPA receptor blocker, 20 uM of NBQX (Cat# 0373, Tocris) was injected to TRN of the aCTC slice following the same described procedures. The chemical inhibition of CTL6 cells was conducted by a global perfusion of clozapine-n-oxide (CNO, 5uM, Cat# 4936, Tocris), the chemical actuator of the chemogenetic probe, hM4Di (*38*) that was solely expressed in CTL6 of NTSR1-Cre mouse after viral injection.

### Electrophysiology and photoinhibition

Whole-cell recording of cortical layer 4 (L4), CTL6, or MGB cells was performed using a visualized slice setup outfitted with infrared-differential interference contrast optics. Recording pipettes were pulled from borosilicate glass capillary tubes and had tip resistances of 2–5 MΩ when filled with potassium gluconate based intracellular solution (in mM: 117 K-gluconate, 13 KCl, 1.0 MgCl_2_, 0.07 CaCl_2_, 0.1 ethyleneglycol-bis(2-aminoethylether)-N,N,N’,N’-tetra acetic acid, 10.0 4-(2-hydroxyethyl)-1-piperazineethanesulfonic acid, 2.0 Na-ATP, 0.4 Na-GTP, and 0.5% biocytin, pH 7.3) for current-clamp mode. Voltage was clamped at −60 mV or +10 mV to measure either the excitatory or inhibitory currents, respectively, using cesium-based intracellular solution (in mM: 117.0 CsOH, 117.0 gluconic acid, 11.0 CsCl, 1.0 MgCl_2_*6H_2_O, 0.07 CaCl_2_, 11.0 EGTA, 10.0 HEPES). Local field potential (LFP) recordings were done using glass pipette with a broken tip (>5um and <10 um) to only allow passing a current around (>500 pA and <1.4 pA as indicated by the membrane test) under a current clamp mode at gain = 100, Bessel = 4kHz.For LFP signals, the data were filtered offline using Clampfit 10.7 software under Gaussian low pass frequency at 300 Hz as well as filtering out the electrical interference at 60Hz. Multiclamp 700B amplifier and pClamp software (Molecular Devices) were used for data acquisition (20 kHz sampling).

For photoinhibition, the halorhodopsin “eNpHR3.0” probe expressed selectively at either CTL6, L5, or IC-GABAergic cells was activated by illuminating a far yellow light (565 nm) obtained from DC fluorescence illuminator (Prior Lumen 200) and Olympus filter cube (U-MF2, Olympus, Japan) which was implemented with TxRed-4040C-000 [excitation:562/40 nm, dichroic 593 nm long pass, emission: 624/40 nm]. The light was set to shine the whole filed of the LFP recording chamber using 4X objective for three seconds extending from one second pre-stimulus and two seconds after the onset of IC stimulation. Based on the initial results related to the dynamics of CTL6 cells, the three seconds illumination was chosen to cover the time period one second before the onset of the stimulus.

### Brain network analysis

The method that was reported before (*39*) and figure S2A shows an overview of the brain network analysis. In this work, we propose an unsupervised framework for brain network classification and investigate how to model and describe neuronal networks using deep neural networks (*40*). In our framework, we utilize a network embedding method (*41*) to find effective representations for brain networks. The dataset used was images of the evoked FA signals obtained from IC, MGB, TRN, and AC of the aCTC slice. 100 instances from each of these brain regions were used. Each instance had one of the ON or OFF labels and consisted of 450 consecutive images from the specific region of the brain. The images of aCTC brain slices have a size of 172*130 pixels. The intensity of the pixels could be detected from these images, and the change of the pixel value could be interpreted as the firing of the underlying group of neurons corresponding to the given pixel. By observing the variation of the pixel value over time, we were able to capture the dynamicity of the signal traveling in the brain slice. A sliding window process was used to grab the timeseries slices between a certain interval. Figure S2B indicates two non-overlapping windows and generated networks for visual purposes, though the algorithm adopts a stride of one timestamp to capture the system evolution at a fine-grained level. For each window, the Pearson product-moment correlation coefficient between all pairs of pixels was calculated and if the correlation was higher than a threshold then an edge with weight equal to the correlation was put between the two nodes representing the two pixels. In our unsupervised architecture, the goal was to learn network embeddings such that networks with similar structure lie close to one another in the embedding space. Two mutually exclusive classes ON/OFF indicating the existence/non-existence of the cortical event, respectively, were used to train and test the classifier. We aimed to learn a mapping function Φ: G → R^k^ that embeds a network G into a k-dimensional space. Our approach used encoder-decoder models (autoencoders) which were trained to reconstruct their input in a way that learn useful properties of the data. The encoder mapped the input to some intermediate representation and the decoder attempted to reconstruct the input from this intermediate representation. The first step to run that was extracting the sequences from the brain networks. we choose LSTMs (Long short-term memory) (*42*) for both the encoder and decoder, forming an LSTM autoencoder (*43*). *LSTM* autoencoders use one LSTM to read the input sequence and encode it to a fixed dimensional vector, and then use another LSTM to decode the output sequence from the vector. Given the trained encoder *LSTM_enc_*, we defined the network embedding function Φ(G) as the mean of the vector’s output by the encoder for network sequences extracted from G:

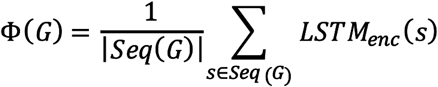

where Seq(G) is the set of sequences extracted from G. Then, Φ(G) is used as the representation of the network G.

We used random walks to generate sequences from brain networks. These sequences are then used to train our LSTM autoencoders. Given a network and a starting node, we selected a neighbor of it at random, and move to this neighbor; then we select a neighbor of this point at random, and move to it etc. The random sequence of nodes selected in this way is a random walk on the network. Given a source node *u*, we generate a random walk *w_u_* with fixed length *m*. Let *v_i_* denote the *i^th^* node in *w_u_*, starting with *v_0_=u*. Then, *v_t+1_* is a node from the neighbors of *v_t_* that was selected with probability 1/d(*v_t_*), where d(*v_t_*) is the degree of *vt*. Figure S2C shows several extracted sequences from a brain network. Each node has an identification number according to the position of the pixel at the brain image. Further, we formulated network representation learning as training an autoencoder on node sequences generated from networks. These autoencoders were based on the sequence-to-sequence learning framework (*44*) [7], an LSTM-based architecture in which both the inputs and outputs are sequences of variable length. The architecture used one LSTM as the encoder *LSTM_enc_* and another LSTM as the decoder *LSTM_dec_*. An input sequence with length *m* is given to *LSTM_enc_* and its elements were processed one per time step. The hidden vector *h_m_* at the last time step m is the fixed-length representation of the input sequence. This vector is provided as the initial vector to *LSTM_dec_* to generate the output sequence. We used the sequence-to-sequence learning framework for autoencoding by using the same sequence for both the input and output. We trained the autoencoder such that the decoder *LSTM_dec_* reconstructs the input using the final hidden vector from *LSTM_enc_* (Figure S2D). We trained a single autoencoder for each region of the brain. The autoencoder is trained on a training set of network sequences pooled across all networks in a region of the brain. After training the autoencoder, we obtain the representation Φ(G) for a single network G by encoding its sequences s ∈ Seq(G) using *LSTM_enc_*, then averaging its encoding vectors. We used (*h_t_*)^*enc*^ to denote the hidden vector at time step t in *LSTM_enc_*, and (ht)^dec^ to denote the hidden vector at time step t in *LSTM_dec_*:

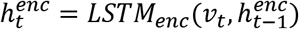

Where *v_t_* is the t^th^ node in a sequence. The hidden vector at the last time step (*h_last_*)^*enc*^ ∈ *R^d^* denotes the representation of the input sequence, and was used as the hidden vector of the decoder at its first-time step:

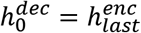

The last cell vector of the encoder was copied over an analogous way. Then each decoder hidden vector (*h_t_*)^*dec*^ is computed based on the hidden vector and node from the previous time step:

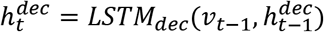

The decoder used (*h_t_*)^*dec*^ to predict the next node vt. For evaluation, we used SVM classifier to classify the brain network representations obtained from our approach (*45*). The 10-fold crossvalidation technique was utilized for the training and test purposes. At each iteration nine folds are used as training data and one-fold as the test case. The average accuracy on the test folds is reported as the final accuracy.

### Classification of the activation state of pre-stimulus activity of CTL6 cells

One second of the pre-stimulus activity of CT-L6 was collected under current clamp modes. The data were down-sampled from 20kHz to 1KHz simplification by using Calmpfit 10.7 software offline. The one-second data set contained 1000 samples indexed in milliseconds for each stimulation. A total of 314 stimulations of IC across 17 cells obtained from 4 brain slices across three animals provided a balanced incidence between ON (55.09%) vs OFF (44.91%) cortical response. Before model training, the dataset underwent preprocessing steps. In order to analyze only the physiologically related frequencies, the frequencies between 0Hz and 50Hz were kept. The filtering procedure consisted of a low-pass filter of order 4. In this way, the filter could correctly remove the frequencies over 50Hz while maintaining the closest phase-shift possible. The Python library for time series feature extraction tsfresh (*46*) was used to process the dataset extracting time series features that were used for the subsequent analysis, including the first 50 components of the Fast Fourier Transform (FFT) (*47*), parameters of a fitted linear regression, entropy (represents the amount of regularity or the level of disorder of the time series), autocorrelation variance and mean, as well as the simple statistics of the time series such as mean, variance and standard deviation. Feature selection (*48*) was performed to select a subset of most informative features. We removed features whose values did not change across all observations. Using a threshold value of 0.6, the cross-correlation was computed for all features to allow the removal of the intercorrelated features that contains the same information for the classification purpose. Finally, the combining scores from ANOVA (*49*) and mutual information (*50*) tests were used to retrieve the 10 most informative features that were used for model training. XGBoost implementation (*51*) of the gradient boosting model (*52*) was used for model training. The binary logistic objective function with a linear booster was chosen as the best fit for the classification task. For model testing, 3-fold cross validation with 10 repetitions was used (*53*). Results were then aggregated over all partitions and cross-validations and the mean and standard deviation was computed. The metrics used to evaluate the model were accuracy and F1-score (*54*). The accuracy is the homogenous measure among all the possible techniques used for the time series classification, thus it will make the results comparable to other possible models. F1 takes into consideration the misclassification error for each class and is explicit about the relative accuracy for each class. The considered baseline is a majority classifier, that is a simple model that always predicts ON class, which has the accuracy equal to 55.09% and F1-score of 71.04% for the ON class and 0% for the OFF class.

### Imaging analysis and statistics

Using customized MATLAB codes, all the pseudocolor images were produced showing the tonotopic map of AC *in-vivo* and the activated brain regions in the aCTC slice. The peaks of FA or Ca signals as well as Δf/f were all computed by Image J software after drawing ROI above the brain region or the individual cell. All the statistical analysis as well as the graphs showing the statistical results were done by Origin-Pro 2017 software. Tests including χ-square, linear correlation fitting, paired t, Repeated measures one-way ANOVA followed by Bonferroni post-test were used across all the work. The significance of the test was set when p-value < 0.05.

### Work art

All figures were designed and made using Adobe illustrator as a part of online Adobe cloud. To keep working within Adobe environment to avoid losing the resolution of the figures, Adobe Photoshop was used to crop the borders of some images to save a space, drawing some scale bars, increasing the darkness and gamma balance of grayscale images showing the electrophysiology recording to be able to show the tiny signals, and writing some titles.

## Results

### Stochastic AC responses to sound presentations *in-vivo*

Consistent with previous work (*55*), the transcranial Ca imaging of the AC of an anesthetized GCaMP6s mouse following repeated presentations of a 5 kHz-37 dB SPL pure tone to the mouse’s right ear (Figure 1A) showed a cortical activity at three distinct areas, the primary AC (A1), secondary AC (A2), and anterior auditory field (AAF) as indicated by sound-evoked Ca-signals shown by MATLAB pseudocolor image (Figure 1B). Such sound-evoked cortical activity (Figure 1B) represented the average of the cortical responses to 10 presentations of the same tone. In contrast, there was schostacity in the cortical responses following the repeated presentations of the same sound as reported by our lab (*20*). Here, the Δf/f of the sound-evoked Ca signals from A1 was computed for every individual trial of sound presentation, and the data showed all-or-none cortical response (Figure 1C). Across the 40 trials of the same sound presentation. the A1 showed a full population response (here called ON cortical responses) or no response (referred to here as OFF responses). ON cortical responses were randomly interrupted by OFF responses (Figure 1C). Collectively, a histogram of Δf/f of sound-evoked Ca-signals of A1 across trials showed two classes of A1 responses (ON vs OFF) (Figure 1D), best represented by a bimodal distribution. (Figure 1E).

**Figure 1:**
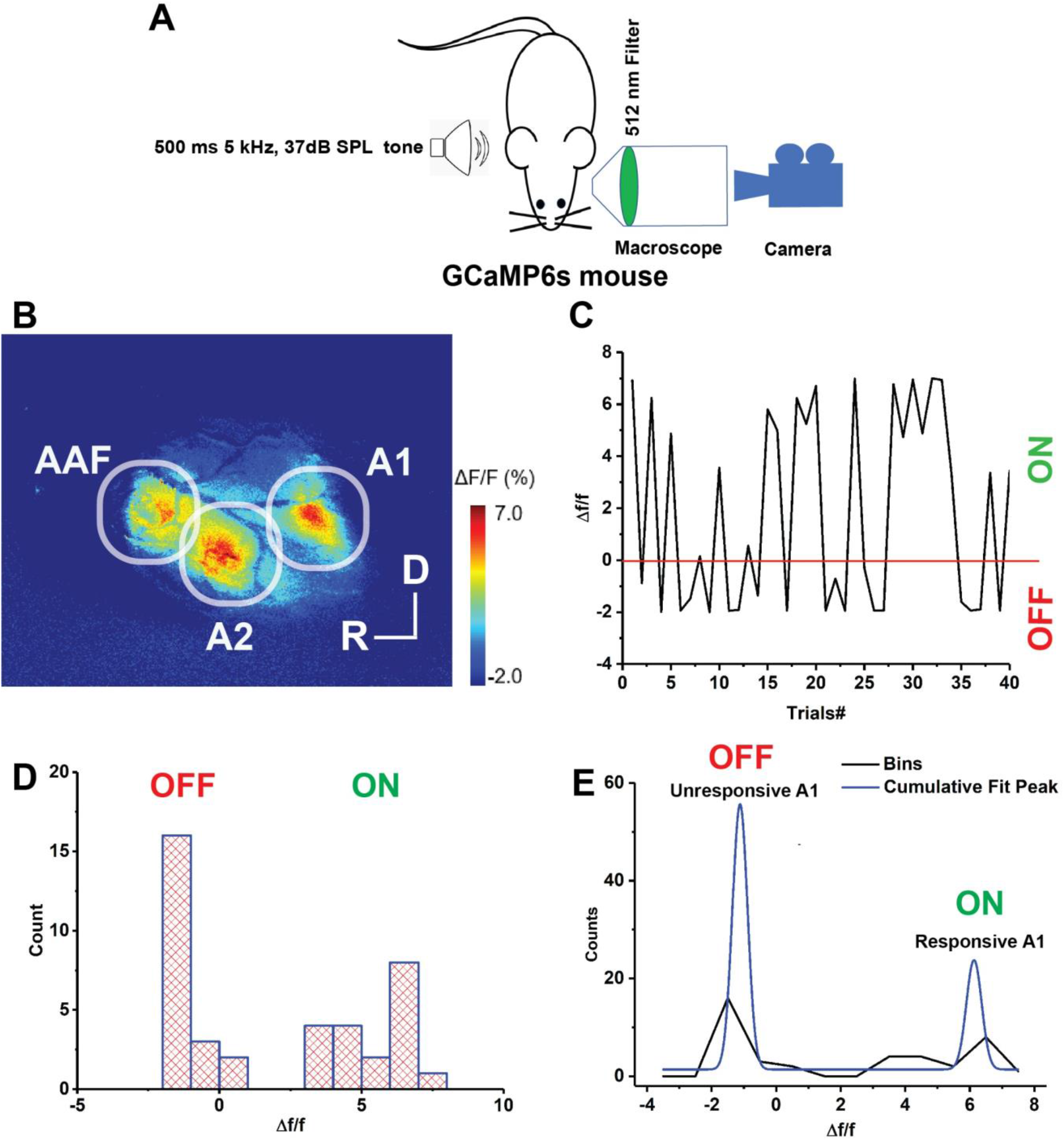
Stochastic auditory cortical population responses to the repeated sound presentations *in-vivo*, A) A cartoon image showing the experimental design of simultaneous transcranial Ca imaging of the left AC of GCaMP6s mouse following playing a pure tone sound at the right ear, B) MATLAB created pseudo-color image representing the average map of AC activation indicated by Δf/f of sound-evoked calcium signals following 10 trials of plying 5kHz, 37 dB SPL pure tone, C) A plot graph of Δf/f of the sound-evoked Ca signals of A1 across 40 trials of plying 5kHz, 37 dB SPL pure tone, D) A histogram of the Δf/f obtained from C, E) A line plot graph of bimodal distribution of two clusters of A1 responses (responsive vs unresponsive A1) [R^2^ = 90.8, ANOVA, f(5,11) = 11.37, p=0.0087], A1: Primary cortex, A2: Secondary cortex, AAF: Anterior auditory filed, D: Dorsal, R: Rostral.

### Stochastic AC responses to electrical stimuli to IC *in-vitro*

To examine the circuit mechanisms underlying the stochasticity of the responses, brain slices that retained connectivity between the IC, MGB, TRN and AC were used. The aCTC mouse brain slice (*31–33*), had an advantage that it retains the synaptic connections between these structures: hence, electrical stimulation of the IC was able to evoke a neuronal activity in all of these brain structures indicated by stimulus-evoked FA and Ca-signals as shown in Figure 2A and B. Both images showed only the average of the stimulus-evoked activity of the connected brain structures in the aCTC slice after several trials of IC stimulation. In contrast, the time series of Δf/f of the stimulus-evoked FA and Ca signals across trials of IC stimulation (Figure 2A and B) showed missing responses at the AC, which was consistent with the *in-vivo* data, despite that MGB, TRN, and IC were always responsive. Looking more closely at the activity of cortical L4 cells, the stimulus-evoked Ca signals of some L4 cells following the IC stimulation showed that each L4 cell had its own profile of responsiveness across trials, but all ceased to respond at trial # 9 and 11, for instance (Figure 2C). To ensure that metabolic or imaging artifact did not drive the observation of these missing cortical responses, whole cell recording of L4 cells as well as LFP recording of L3/4 were conducted. During the OFF cortical responses indicated by the absence of FA signals, L4 cells did not fire action potentials (Figure 2D), and similarly, there were no LFP signals recorded from cortical L3/4 (Figure 2E), which implicated that FA imaging reflected the real status of the cortical activity following the stimulus presentation, and the missing cortical responses represented a lack of signal propagation within the cortex. Interestingly, during ON cortical responses, the IC stimulation evoked a cortical UP state in L4 cells, which was associated with many action potentials (Figure 2D, red box). Consistent with that, LFP recordings showed post-stimulus strong responses during ON cortical responses following the stimulation of the IC as reported before (*56*) (Figure 2E, green box). Stochastic AC responses were also seen in brain slices prepared using a different anesthetic (isoflurane) without trans cardiac perfusion and in a laboratory with a biphasic stimulator vs. monophasic stimulator. The results in this case were identical (figure S3), suggesting that the stochastic responses were not specific to a particular test preparation.

**Figure 2:**
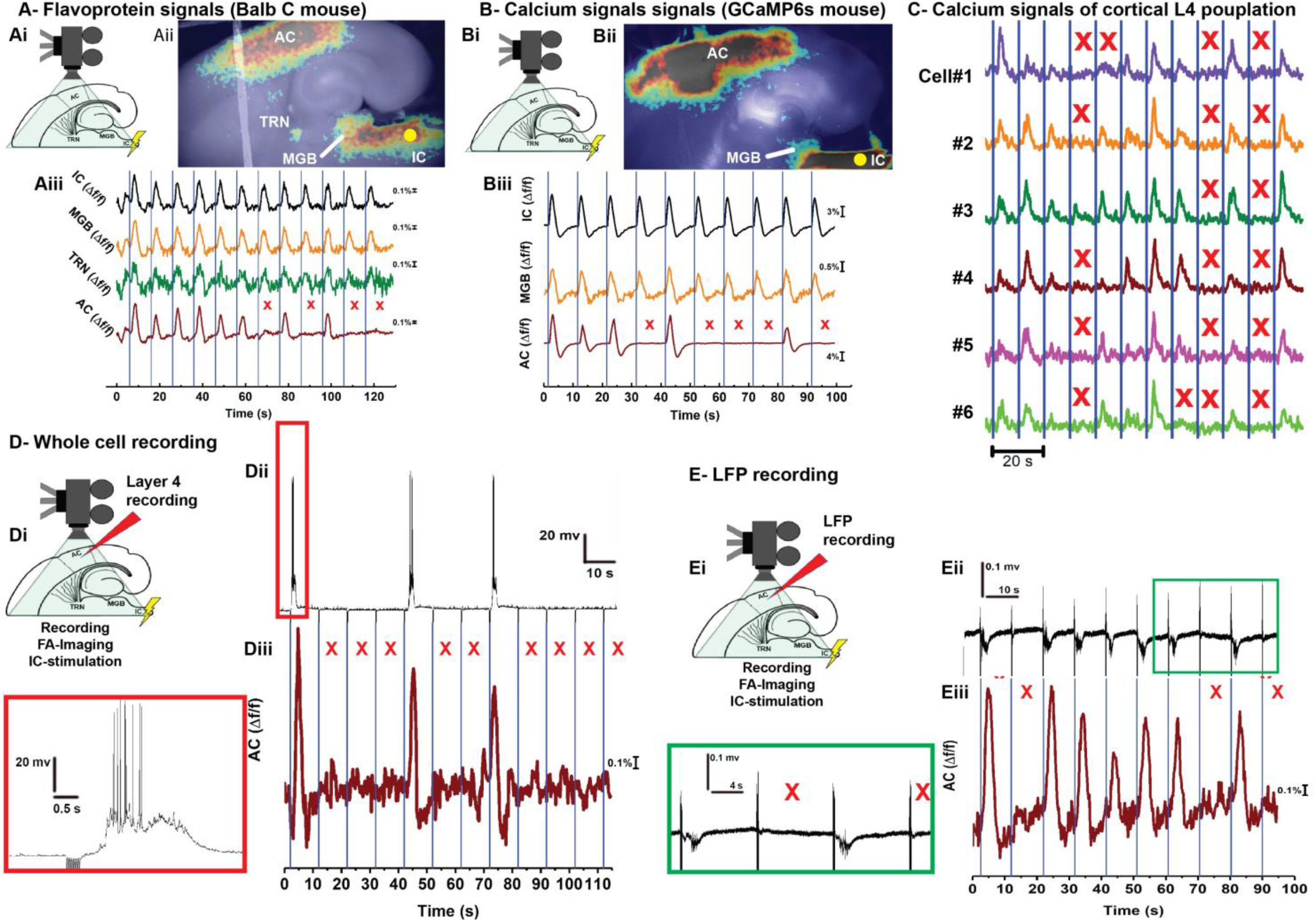
Stochastic auditory cortical responses to the repeated sound presentations *in-vitro:* A-B) Ai and Bi: Cartoon images showing the experimental design of simultaneous FA or Ca imaging, respectively, and the IC stimulation of the aCTC slice, Aii and Bii: MATLAB pseudo-color images showing the neuronal activation indicated by evoked FA or Ca signals, respectively, in the IC, MGB, TRN and AC following the IC stimulation, Aiii and Biii: The time series of Δf/f of evoked FA or Ca signals, respectively, in the IC, MGB, TRN and AC following the IC stimulation, C) The time series of evoked Ca signals of some activated L4 cells following the IC stimulation, D) Di and Ei: Cartoon images showing the experimental design of the simultaneous FA imaging and whole cell recording of L4 cells or LFP recording of layer 3/4, respectively, Dii and Diii: The time series of L4 whole cell recording and the Δf/f of the evoked cortical FA signals, respectively, following the IC stimulation, Eii and Eiii: The time series of the L3/4 LFP signals and Δf/f of the evoked cortical FA signals, respectively, following the IC stimulation, red box: The magnification of the post-stimulus activity of L4 cells showing evoked upstate associated with action potentials, green box: The magnification of some post-stimulus cortical LFP signals showing UP state evoked activity, red Xs refer to the occurrence of OFF cortical responses indicated by the absence of cortical FA or Ca signals as well as post-stimulus cortical LFP signals, blue lines indicate the onset of the IC stimulation, Yellow circle indicates the position of the electrical stimulation of the IC,

OFF cortical responses occurred only after electric stimulation of IC, and not by the direct stimulation of MGB or the subcortical white matter (Figure S3). According to the observation that the OFF responses were shown only by the AC despite the activation of the other subcortical brain structures of the circuit in the aCTC slice, the images of the evoked FA signals at the subcortical structures (IC, MGB, and TRN) following the IC stimulation were analyzed using a deep learning algorithm (*41*) to determine which subcortical structure could best predict whether a cortical response would be an ON or an OFF response based on a trained classifier SVM (*45*). Figure S2E shows a few examples of the correlation networks constructed from our data. The blue points are the network’s nodes that represented the activated pixels of the image indicated by evoked FA signals following the IC stimulation. These representations were used in a brain network classification task using our proposed model on each subcortical structure separately and compared the accuracy of classification on these structures. SVM classifier showed that the activated pixels of MGB following IC stimulation had a higher accuracy (81%) for the classification between ON and OFF cortical responses compared to 78% for IC and 74% for TRN. Such finding suggested that MGB could play a critical rule in modulating the cortical activity.

### The OFF cortical responses are driven by inhibition in the MGB

Since the OFF cortical response represented a full absence of stimulus-evoked cortical activity, we reasoned that OFF cortical responses could be driven by inhibition. To do that, the global disinhibition in the aCTC slice by bath application of gabazine, the GABAα-R blocker, was conducted. Under simultaneous FA imaging and IC stimulation, gabazine perfusion was able to retrieve all missing cortical responses compared to control and washing indicated by the Δf/f of evoked cortical FA signals (Figure 3A), which suggested that the OFF cortical responses were driven by inhibitory inputs. The AC and MGB were further investigated to search for the site of inhibition that drove such OFF cortical responses. Rationally, observing evoked post-stimulus IPSCs from any of these brain structures during the OFF cortical responses could lead to uncovering the site of inhibition. As such, the whole cell recording of cortical L4 or MGB cells at +10 mV voltage clamp was conducted simultaneously with FA imaging following the stimulation of the IC to track the IPSCs in two cell types during ON vs OFF cortical responses. Although L4 cells demonstrated a surge of evoked post-stimulus IPSCs during ON cortical responses, they did not show any evoked post-stimulus IPSCs during OFF cortical responses which suggested that cortex did not receive inhibitory signals during OFF cortical responses (Figure 3B). In contrast, MGB cells showed evoked post-stimulus IPSCs following every trial of the IC stimulation during ON and OFF cortical responses (Figure 3B), which was consistent with the fact that MGB is always active after each trial of IC stimulation (Figure 2). However, the inhibitory and excitatory currents received by MGB cells were further analyzed to investigate any change in both currents during ON vs OFF cortical responses. Interestingly, the evoked post-stimulus IPSCs in the MGB cells were larger during OFF compared to ON cortical responses with no difference in the net excitatory transferred charges (Figure 3C). This finding suggests that MGB cells receive more inhibition during OFF cortical responses with no change in excitation, which led us to hypothesize that MGB activity could be modulated by inhibitory inputs during the OFF cortical responses. To test this hypothesis, the disinhibition in the MGB was examined to determine if it can retrieve the missing cortical responses. Under simultaneous FA imaging and IC stimulation, the specific injection of gabazine into MGB nucleus in the aCTC slice was able to retrieve the missing cortical FA signals indicated by the Δf/f of the evoked cortical FA signals (Figure 3D). In contrast, the selective gabazine injection into the AC was not able to retrieve the missing cortical responses (Figure 3D), which was consistent with the outcome of electrophysiology data obtained from the whole cell recording of L4 cells (Figure 3B). Accordingly, these data confirmed that the OFF cortical responses could be driven by thalamic inhibition.

**Figure 3:**
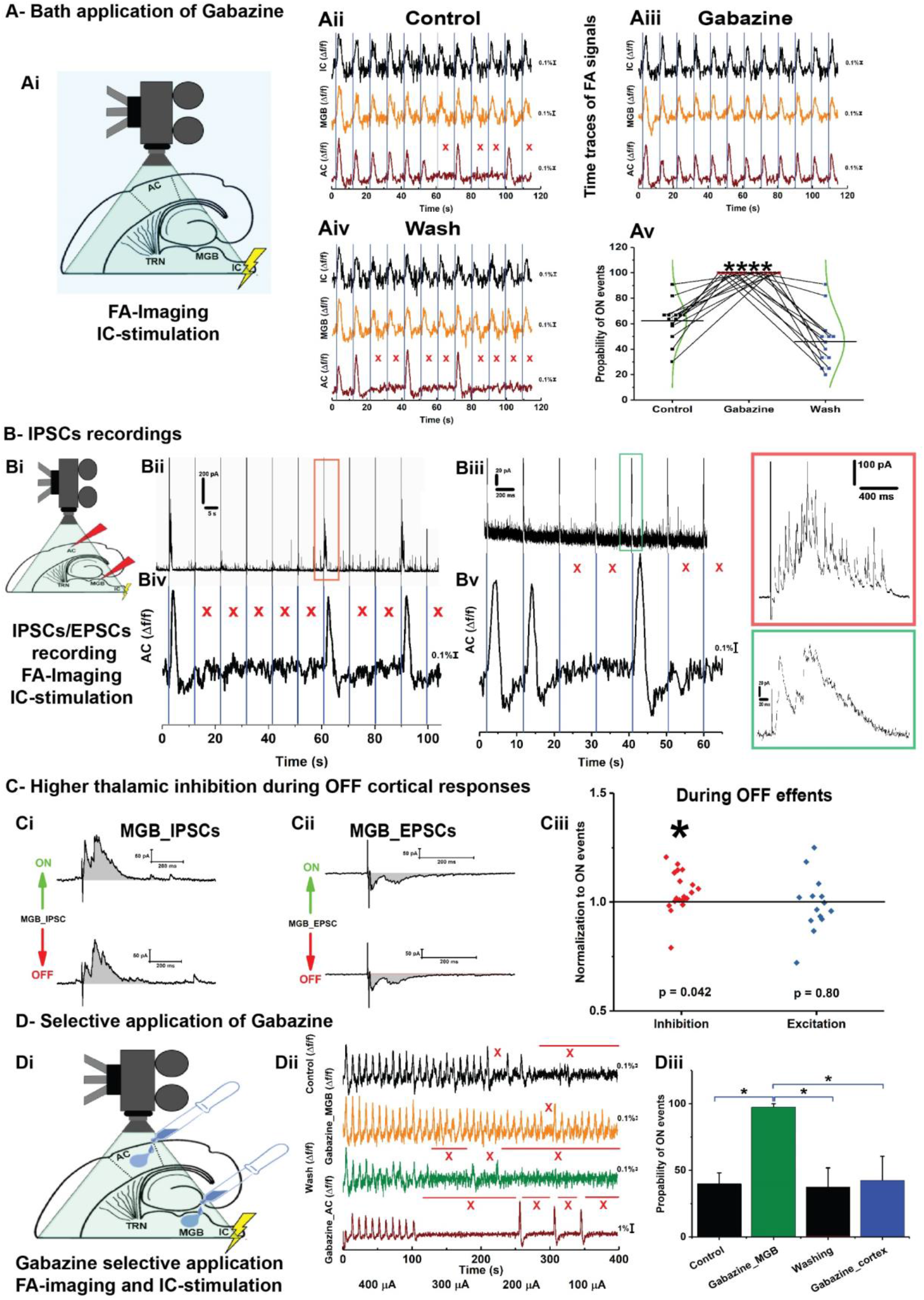
The OFF cortical responses are driven by MGB inhibition. A) Ai: A cartoon image showing the experimental design of simultaneous FA imaging and IC stimulation, Aii-Aiv: The time series of Δf/f of the evoked cortical FA signals in the IC (black trace), MGB (orange trace), and AC (brown trace) following the stimulation of the IC under ACSF (control), (gabazine), or washing by ACSF (Wash), respectively, Av: A plot graph of the results of repeated measures one-way ANOVA showing that the probability of ON cortical responses was significantly higher than that of control and wash [**** f(1,12) = 731.16, p < 10^−9^, p =3.3×10^−6^ for control vs gabazine, p = 0.027 for control vs wash, p < 10^−9^ for gabazine vs wash], B) Bi: A cartoon image showing the experimental design of simultaneous FA imaging, IC stimulation, and IPSCs or EPSCs recording from L4 or MGB cells, Bii-Biii: Evoked post-stimulus IPSCs recorded from L4 or MGB cell, respectively, Biv-Bv: The time series of Δf/f of the evoked cortical FA signals simultaneously imaged during IPSCs recording from L4 or MGB cell, respectively, red box and green box: The magnification of the evoked post-stimulus IPSCs recorded from L4 or MGB cell, respectively, C) Ci-Cii: Evoked post-stimulus IPSCs or EPSCs, respectively, recorded from MGB cell during ON vs OFF cortical events, Ciii: A scatter plot graph of the area under the curve (AUC) of the evoked post-stimulus IPSCs and EPSCs recorded during OFF cortical responses normalized against those recorded during ON cortical events showing that MGB cells received more inhibitory charges during OFF cortical events with no change in the received excitatory charges [paired t-test: for EPSCs, *t(13) = 0.26, p = 0. 8, for IPSCs, t(18) = −2.18, p = 0.042], D) Di: A cartoon image showing the experimental design of simultaneous FA imaging, IC stimulation, and selective gabazine injection into MGB or AC using picospritzer, Dii: The time series of Δf/f of the evoked cortical FA signals during the injection of ACSF (control, top black trace), gabazine into MGB (orange trace), gabazine into AC (brown trace), or after washing (middle green trace), Diii: A bar graph of repeated measures one way ANOVA results showing that the probability of ON cortical responses were significantly higher after the injection of gabazine into MGB [*f(1,3) = 35.76, p = 0.009, p = 0.01 for control vs gabazine into MGB, p = 1 for control vs gabazine into AC, p = 1 for control vs wash, p = 0.016 for gabazine into MGB vs gabazine into AC, p = 0.009 for gabazine into MGB vs wash, p = 1 for gabazine into AC vs wash], red Xs refer to the occurrence of OFF cortical responses indicated by the absence of cortical FA or Ca signals as well as post-stimulus cortical LFP signals, blue lines indicate the onset of the IC stimulation, AC: IPSCs: Inhibitory postsynaptic currents, EPSCs: Excitatory postsynaptic currents.

### CTL6 cells are the main driver of the missing cortical response via TRN

Further work was done to investigate from where MGB received these inhibitory inputs to drive the cortical OFF responses. As previously reported, MGB can be inhibited mainly by IC through the feedforward inhibition by IC GABAergic cells (*57, 58*) or by the cortex through the feedback inhibition by TRN (*59*). As such, the disinhibition of these pathways was examined to determine if it can retrieve the missing cortical responses. For inhibition of the feedforward inhibition, the IC of neonatal GAD2-Cre mice was injected with halorhodopsin-AAV virus (See Methods) to induce the expression of eNpHR3.0 or halorhodopsin specifically in the GABAergic cells of the IC in a Cre-dependent manner. As expected, GABAergic cells of the IC as well as their projections to MGB expressed halorhodopsin indicated by the presence of YFP (Figure S5A). The photoinhibition of GABAergic cells of the IC by full field illumination of 565 nm light was not able to retrieve the missing cortical responses indicated by no recovery of the post-stimulus LFP signals recorded from L3/4 (Figure S5B). The statistical analysis showed no significant difference in the probability of ON cortical responses with and without the photo-inhibiting light (Figure S5C), which suggested that the OFF cortical responses were not driven by the feedforward inhibition of MGB. CTL6 cells also send inhibitory signals to thalamic cells through indirect inhibitory synapses via TRN, a shell-like structure of GABAergic neurons that surrounds the most of dorsolateral part of the thalamus (*60–63*). Therefore, the feedback inhibition of MGB via CTL6-TRN pathway was examined. The injection of the AC of NTRS1-Cre neonates with Ha-AAV1 resulted in a successful Cre-dependent expression of eNpHR3.0 receptors in the CTL6 as well as their projections to TRN and MGB (Figure 4A). Interestingly, the photoinhibition of CTL6 cells by full field illumination of 565 nm light resulted in a significant increase of the probability of ON cortical responses as indicated by the recovery of the post-stimulus LFP signals from L3/4 compared to the control (No light) (Figure 4A). To further prove the previous outcome, the AC of a separate group of NTRS1-Cre neonates was injected with DREADDs-AAV virus, which resulted in a successful Cre-dependent expression of the inhibitory chemogenetic receptors, hM4Di, as indicated by m-cherry tag specifically in CTL6 cells as well as their projections to TRN and MGB (Figure S6A). Consistent with the previous data, the chemical inhibition of CTL6 cells expressing hM4Di receptors by their chemical actuator, CNO, significantly increased the frequency of ON cortical responses as indicated by Δf/f of evoked cortical FA signals compared to the control (Figure S6B and C). Given that CTL6 cells project to MGB through a direct excitatory synapse, further examination was required to test if the TRN is the main driver of CTL6 effect. Blocking of the TRN activity by the NBQX, the AMPA-R blocker (*36, 37*), which was specifically injected into TRN, significantly increased the probability of ON cortical events indicated by Δf/f of the evoked cortical FA signals (Figure 4B). Accordingly, based on the data shown in figure 4, we conclude that the OFF cortical responses were driven by the feedback inhibition of MGB by CTL6 cells via TRN.

**Figure 4:**
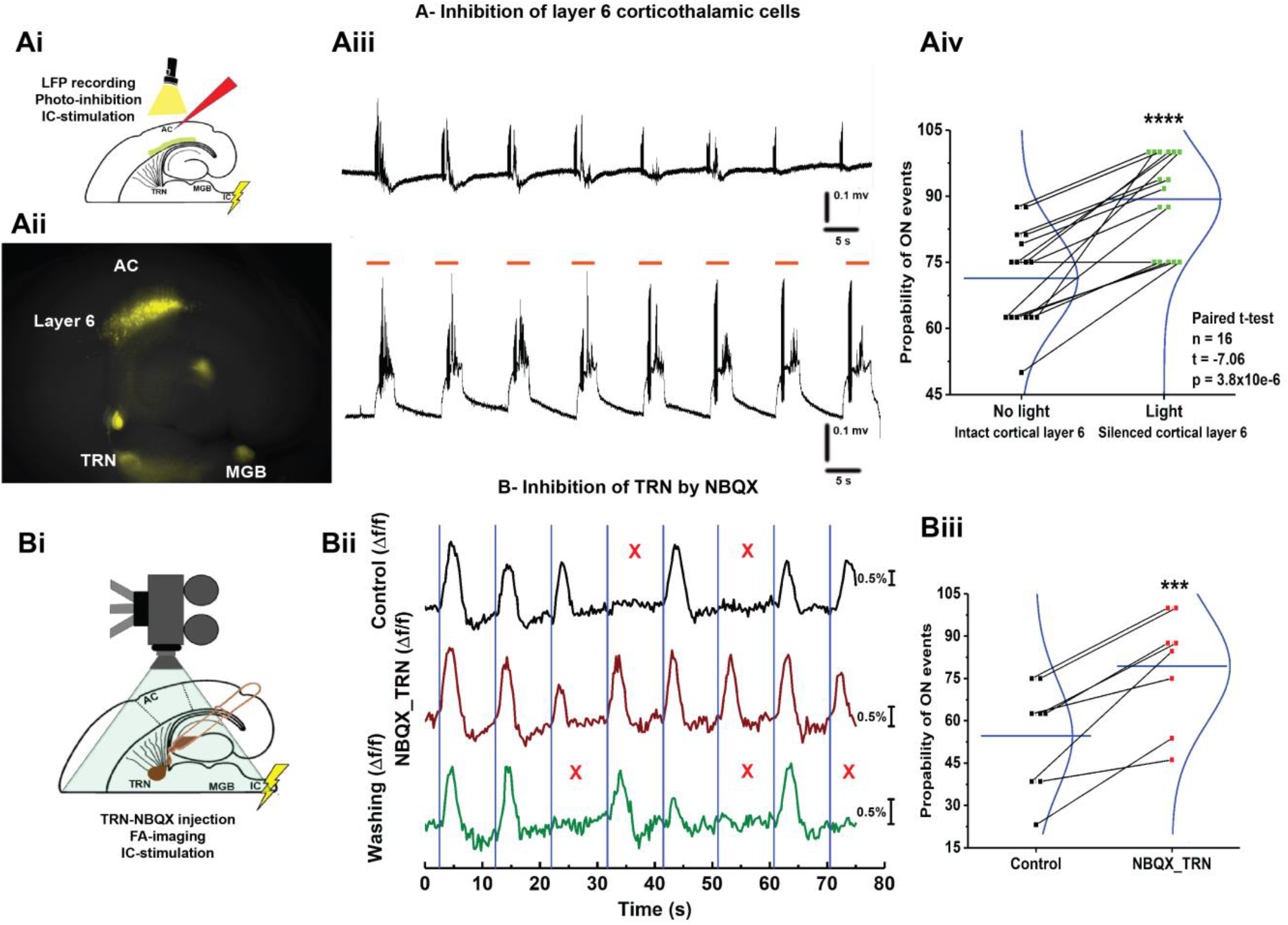
The OFF cortical responses were driven by the feedback inhibition of MGB by CT-L6-TRN pathway: A) Ai: A cartoon image showing the experimental design of simultaneous IC stimulation, LFP recording, and full field photoinhibition, Aii: Image of aCTC slice of NTSR1-Cre mouse showing the expression of eNpHR3.0 receptors as indicated by YFP tag in NTSR1 +ve CTL6 cells as well as their projections to TRN and MGB, Aiii: The time series of the post stimulus cortical LFP signals from L3/4 following the IC stimulation without (top) and with 565 nm light (bottom), Aiv: A scatter plot graph of paired t-test showing that the probability of ON cortical events was higher during the photoinhibition of CTL6 cells by 565 nm light [paired t-test, ****t(15) = −7.06, p = 3.8X10^−6^], B) Bi: A cartoon image showing the experimental design of simultaneous IC stimulation, FA imaging and NBQX injection into TRN by Picospritzing, Bii: The time series of Δf/f of the evoked cortical FA signals during ACSF (black trace) and NBQX (brown trace) injections into TRN as well as washing (green trace), right panel, Biii: A plot graph of paired t-test showing that the probability of ON cortical events was higher by blocking Trn activity by NBQX compared to the control [paired t-test, ***t(5) = −6.03, p = 5.2X10^−4^], red Xs refer to the occurrence of OFF cortical responses indicated by the absence of the post-stimulus LFP signals recorded from L3/4 or Δf/f of the evoked cortical FA signals, orange lines indicate the time period of illuminating the light, blue lines indicate the onset of the IC stimulation.

However, it is not clear how these random OFF cortical responses could be driven under the control of CTL6 cells. Given that intertrial variability of sensory-evoked cortical responses could be dependent on the state of intrinsic cortical activity which is in turn stochastic in nature (*4, 10, 64, 65*), the oscillation of the spontaneous activity of the CTL6, one second before the stimulus onset, was examined to determine if it can be used to build a classifier that predicts the cortical responses (ON vs OFF). Initially, a time period of one second before the stimulus onset was taken from the time trace of membrane potential recording of CTL6 cells during ON vs OFF cortical responses (Figure S7B). These time periods were used to encode and train the classifier, then other time periods were used for testing (See methods). Table S1 shows the results computed using the proposed classification method on the considered dataset, and the confusion matrix showed the results obtained from the classifier (Figure S7C). Interestingly, the one second pre-stimulus activity of CTL6 cells (≤ 50 Hz frequency) was 63.06% accurate to predict the cortical response (ON vs OFF). Such prediction was significantly higher than the baseline accuracy (55.09%) (Figure S7F). The proposed method was able to provide a better classification than the majority classifier (9% above baseline prediction). These data suggest that the oscillation of CTL6 cells could modulate MGB activity to gate the flow of the sensory information to control the cortical sensory gain. To test the specificity of CTL6-TRN pathway to modulate MGB activity, a negative control experiment was done by inhibiting L5 cells that do not have any projections to TRN (*66*). The injection of the AC of RPB4-Cre neonatal mice with Ha-AAV1 virus resulted in a successful Cre-dependent expression of eNpHR3.0 receptors in L5 cells as indicated by YFP tag (Figure S8B). Consistent with the previous data, the photoinhibition of L5 cells expressing eNpHR3.0 receptors by full field illumination of 565 nm light could not retrieve the missing cortical responses indicated by no recovery of the post-stimulus LFP signals from L3/4 compared to control (No light) (Figure S8B and C), which implicated that the OFF cortical responses were specifically driven by MGB inhibition through CTL6-TRN pathway.

### Synchronized MGB cells are associated with ON-cortical responses

Since MGB was always activated following each trial of IC stimulation, how could inhibition modulate its activity? To answer this question, we hypothesized that despite the evoked thalamic activity, thalamic cells can only evoke a cortical response under a prerequisite spatial, temporal, or spatiotemporal coordination. To test this hypothesis, Ca signals of MGB cells were imaged following the stimulation of the IC to examine if there was a change in the spatial and/or temporal activation of MGB cells during ON vs OFF cortical response (Figure 5A). Our data showed that the evoked Ca signals demonstrated three categories of activated thalamic cells. The first and second categories (Cat. #1 and 2) represented thalamic cells that were exclusively activated during ON or OFF cortical responses (Figure 5B, green dots for ON and red dots for OFF). However, the last category represented the thalamic cells that were activated during ON and OFF cortical responses without preference, so they had no spatial difference (Figure 5B, yellow dots). Further, the variance of the peak latencies of the evoked Ca signals from all thalamic cells in the three categories was calculated to examine the temporal difference between cells. Interestingly, Cat# 1 and 2, which were spatially different, showed no difference in the variance of the peak latencies of their evoked Ca signals, which indicated no temporal difference (Figure 5C). In contrast, category 3 MGB cells showed a higher variance of their peak latency during OFF cortical response (Figure 5E and F), which suggested that this population of MGB cells had unsynchronized activation during OFF cortical responses. To visually indicate this temporal difference in the activation of category 3 MGB cells during ON vs OFF, the histogram demonstrated that there were more synchronized MGB cells during ON cortical response indicated by the small deviation of the peak latency of their Ca signals around the mean (Figure 5G, green arrow), while the activated MGB cells during OFF cortical responses had an increased spread (Figure 5G, red arrows). These data suggest that synchronous thalamic relay cell activity is required to evoke a cortical ON response, as has previously been suggested (*67*).

**Figure 5:**
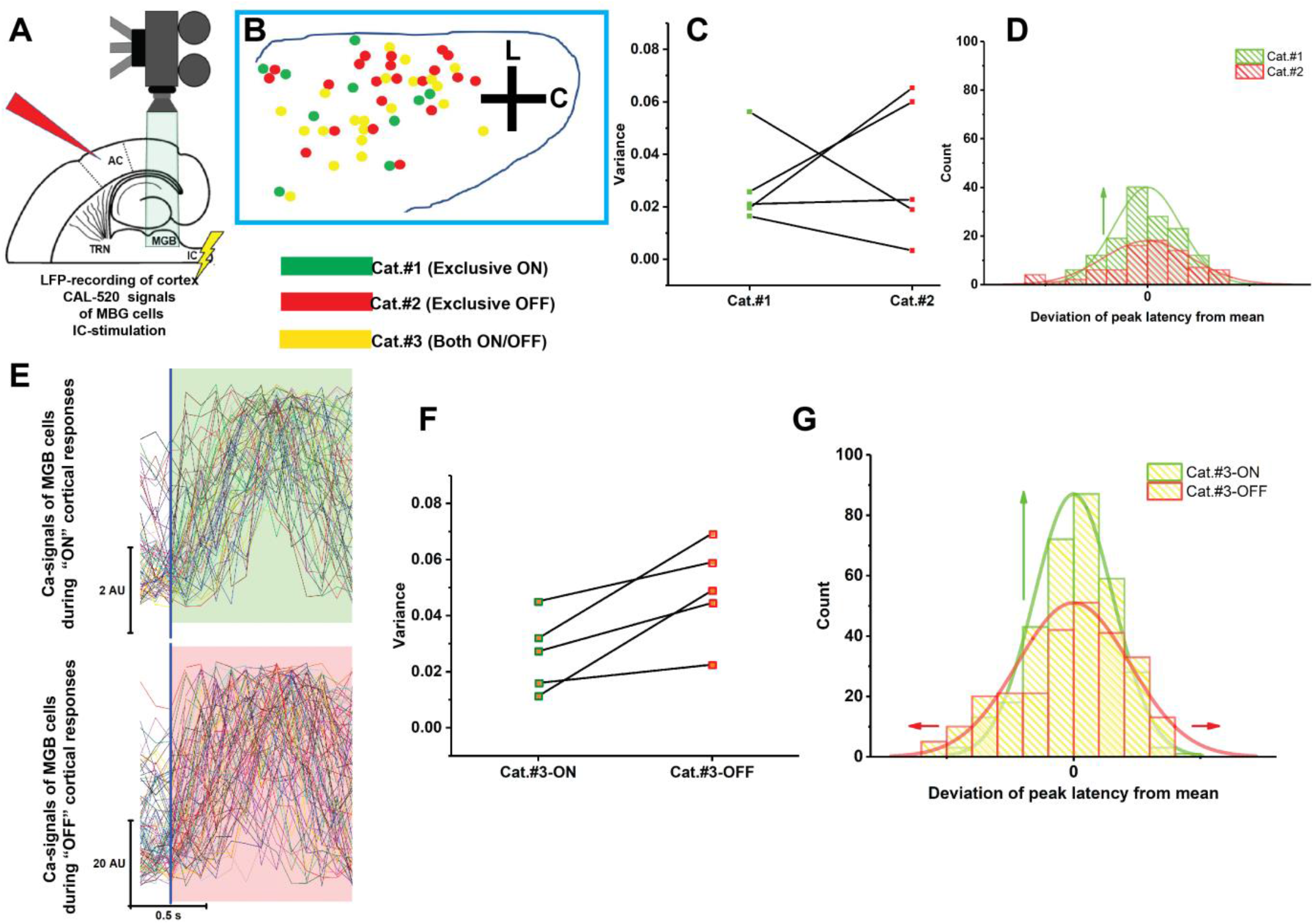
Unsynchronized MGB cellular activity is associated with OFF-cortical responses. A) A cartoon image showing the experimental design of simultaneous IC stimulation, Ca imaging of MGB cells, and cortical LFP recording, B) A cartoon image showing the locations of the activated thalamic cell, indicated by evoked Ca signals following IC stimulation, during ON only (green circles), OFF only (red circles), and during both (yellow circles), C) A scatter plot graph of the paired t-test showing no change between the variances of peak latencies of cat.#1 and 2, [t(4)= −0.41, p = 0.71], D) A histogram of the deviations of peak latencies from their mean of the Ca signals of Cat.#1 cells (green bars and line) vs Cat.#2 cells (Red bars and line), E) The sweeps of the evoked Ca signals of all activated cells during ON (top) vs OFF (bottom) cortical responses indicated by the post stimulus cortical LFP signals recorded from L3/4, F) A scatter plot graph of the paired t-test showing a higher variance of the peak latencies of Cat.#3 cells activated during OFF cortical response compared to those activated during ON cortical responses [*t(4) = −3.56, p = 0.024], F) A histogram of the deviations of the peak latencies from their mean of the Ca signals of Cat.#3 cells activated during ON (yellow bars and green line) vs OFF cortical responses (yellow bars and red lines).

## Discussion

We observed stochastic population cortical responses in the mouse AC following exposure to pure tones *in-vivo* or electrical stimulation of the IC *in vitro*. Population ON responses were associated with preceding oscillations in layer 6 corticothalamic neurons and with synchronized responses among MGB cells. Population OFF responses were associated with TRN-mediated inhibition at the level of the MGB, under the control of layer 6 corticothalamic projections. Other inhibitory projections to the MGB from the IC had no impact on the probability of eliciting an ON response. Below, we discuss these data in the context of sensory processing in thalamocortical systems.

During ON cortical responses, the TC afferents evoked an UP state in L3/4 consistent with that shown in murine TC brain slice (*34*). These TC evoked UP states resembled the cortical UP states reported *in vivo* and *in vitro* initiated spontaneously by intracortical networks (*15, 68, 69*) or by TC inputs (*2, 56*). Given that the UP states evoked spontaneously and by TC inputs shared the defined temporal sequence, MacLean *et al*. (*2*) suggested that the predefined cortical circuits may govern and dominate the TC inputs as described previously (*9*). Supporting this idea, the OFF cortical responses were observed in the AC despite the activation of IC, MGB, and TRN following IC stimulation, contradicting expectations of a linear filter model that cortex should respond as long as MGB linearly transmits information upon its activation. Also, OFF cortical responses were not a sign of cortical adaptation which is characterized by gradual decrease of the response due to the repeated presentation of the stimulus (*70, 71*). Previous work has shown that sensory-evoked cortical variability was predictable based on the magnitude and the phase of pre-stimulus ongoing cortical oscillations (*72, 73*). Such a relationship means that the state of the cortical network could shape the sensory-evoked cortical responses. In fact, CT axons are about 10:1 greater than TC ones (*74, 75*) and more than 40% of the synapses thalamic cells are formed by CT projections (*76–78*), which suggests that cortex has a strong influence on thalamic activities. In fact, upon activation by CTL6, TRN was reported to send inhibitory inputs to thalamic relay cells that could modulate thalamic activity (*63, 79–81*). Interestingly, the photo-stimulation of cortical L6 was reported to reduce the visually evoked activity in LGN relay neurons (*25*). Consistent with that finding, our data demonstrated that both photo-inhibition of CTL6 cells and blockade of TRN activity were able to retrieve the missing cortical responses, thus implicating di-synaptic feedback inhibition of MGB by CTL6 cells was the main control point of the OFF cortical responses. MGB inhibition induced by the TRN could modulate the spatiotemporal coordination between thalamic cells due to the heterogeneity of TRN cells for their intrinsic properties and their axonal arbor size (*82, 83*), which supported our finding that there was a specific spatiotemporal coordination between MGB cells exclusively during ON vs OFF cortical responses (Figure 5).

In addition to the indirect inhibitory projections to MGB from CTL6 via TRN, there are also direct excitatory projections from CTL6 cells to MGB (*59, 84–86*). In the TC somatosensory slice, CTL6 cells can bidirectionally switch their excitability to favor the activation or the suppression of the somatosensory thalamus depending on the oscillation of their evoked activity (*63*). *In-vivo* whole cell recording of L2/3 cells in V1 showed that the visually evoked 3-5 Hz membrane potential reduces the responsiveness of the visual cortex (*87*), which suggests that the internal dynamics of cortical CTL6 cells could alter evoked cortical activity. OFF cortical responses in the current study always occurred after a single or multiple IC stimulation, which could suggest that OFF cortical responses occurred only after evoked cortical activity that could change the internal dynamics of the cortical cells.

### Experimental considerations

Since the *in vivo* experiments were done here on anesthetized mice, it is not known whether ON- or OFF-responses would correlate to behavioral responses in an awake animal. However, the presence of ON- and OFF-responses in an anesthetized animal does not exclude them as a potential substrate for perception, since the anesthesia may prevent behavioral responses through other mechanisms (just as sensory cortical responses are preserved during sleep (*88, 89*)). In addition, intertrial variability of sensory-evoked cortical responses has been observed in awake and anaesthetized animals (*73, 90, 91*). Further, in the auditory system, it was found that primary AC individual cells maintained the strengths of their evoked activity to pure tones in both awake and ketamine/xylazine anesthetized conditions (*92*). Forthcoming work will determine the relationship between ON- and OFF-cortical responses and conscious perception of threshold stimuli or stimuli in noise.

### Implications

Previous studies viewed the thalamus and cortex as a series of filters whereby combinations of receptive fields produce increasingly selective feature detectors, culminating in uniquely selective neurons (i.e., “grandmother cells”). This type of organization implies that moment-to-moment perception is a reflection of detailed streams of information coursing through ascending sensory systems, to be consciously perceived when those streams engage highly selective cells in the cortex. An alternative view is that ascending information is used to create and modify a bank of sensory representations that get called upon depending on behavioral needs, and that conscious perception reflects activation of these pre-wired circuits. The former theory views the thalamocortical synapse as a unit of perception, while the latter views this synapse as a unit of learning.

A growing body of literature supports the notion that conscious perception involves the release of stereotyped patterns of cortical activity. Cortical circuits undergo spontaneous activity that is thought to be the substrate for ongoing thought, memories, and dreams (*11, 12, 16, 17*). Sensory inputs appear to engage the same cortical patterns (*2–5*), suggesting that the role of thalamocortical transmission is to activate cortical ensembles rather than impress sensory information upon them. This view comports with the finding that loss of afferent input, or increased uncertainty about afferent input, leads to untethering of the cortex and subsequent spontaneous patterns of sensory cortical activity (i.e., hallucinations). (*93*).

Critical for any such mechanism of control of cortical ensembles is a means by which those ensembles are selected. The current data suggest that the TRN, under the control of layer 6 corticothalamic projections, activates populations of thalamic neurons by synchronizing their responses, increasing the likelihood of engendering a population cortical response. This mechanism of TRN-based thalamic synchrony to activate the cortex has been proposed previously (*94*) and is consistent with the finding that populations of thalamic neurons are required to optimally activate the cortex (*67*) and that the TRN is at the heart of a prefrontal cortex-based mechanism to shape cortical activation under changing cognitive demands (*95–97*). Further, the TRN receives inputs from basal forebrain, amygdala and non-reciprocally linked regions of the thalamus (*98–105*), forming an assortment of inputs to potentially modulate TRN, and ultimately select cortical circuits. Given the putative role of the TRN in the selection of thalamic, and therefore cortical, circuits during sensory perception, one would predict that disruption of TRN activity could lead to uncontrolled release of patterns of cortical activity. Consistent with this idea, ample evidence has accumulated to suggest that schizophrenia, a disease characterized by the presence of auditory hallucinations, involves disruption of the TRN (*106–115*).

### Conclusion

Here, we described a unique stimulus-evoked population cortical all-or-none response, which suggests that thalamus recruits cortical ensembles of a pre-wired sensory representations upon external stimulation and internal cortical dynamics of corticothalamic neurons. These data also suggest that corticothalamic modulators control the spatiotemporal coordination between the thalamic cells to gate the thalamic ability to activate the intracortical network. It will be important in future studies to more fully understand how other regulators of the TRN, such as the basal forebrain, prefrontal cortex and amygdala, influence the selection of cortical circuits during behavior.

## Acknowledgments

This work was supported by NSF1515587, DC013073 and DC014765

---

**Figure 1S:**
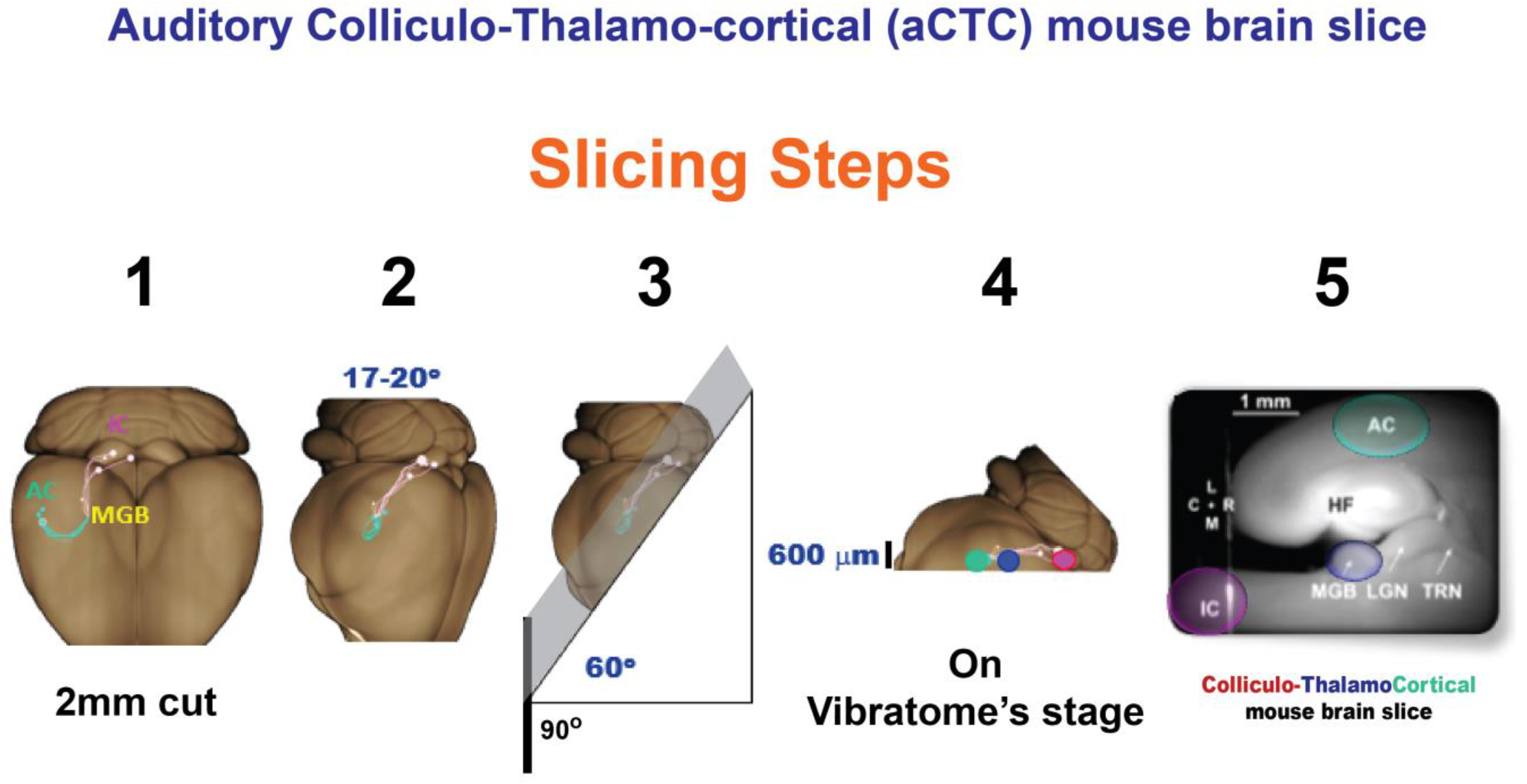
Slicing steps of aCTC mouse brain slice: 1) The brain was cut from its olfactory bulb as 2mm to the caudal axis to make the first platform, 2) The brain was lifted on its frontal aspect on the cut surface made by step 1, then rotated 17-20° angle relative to the brain’s midline, 3) From the dorsal view, the brain was then cut at 60° angle relative to the base and perpendicular on the horizontal line to make the second platform, 4) The brain was lifted on the cut surface made by step 4 on vibratome’s stage and cut as 600 um slices, 5) The final look of aCTC slice should have IC, MGB, and AC connected together at one spatial plan, AC: Auditory cortex, C: Caudal, FA: Flavoprotein, IC: Inferior colliculus, L: Lateral, LGN: Lateral geniculate nucleus, M, Medial.

**Figure 2S:**
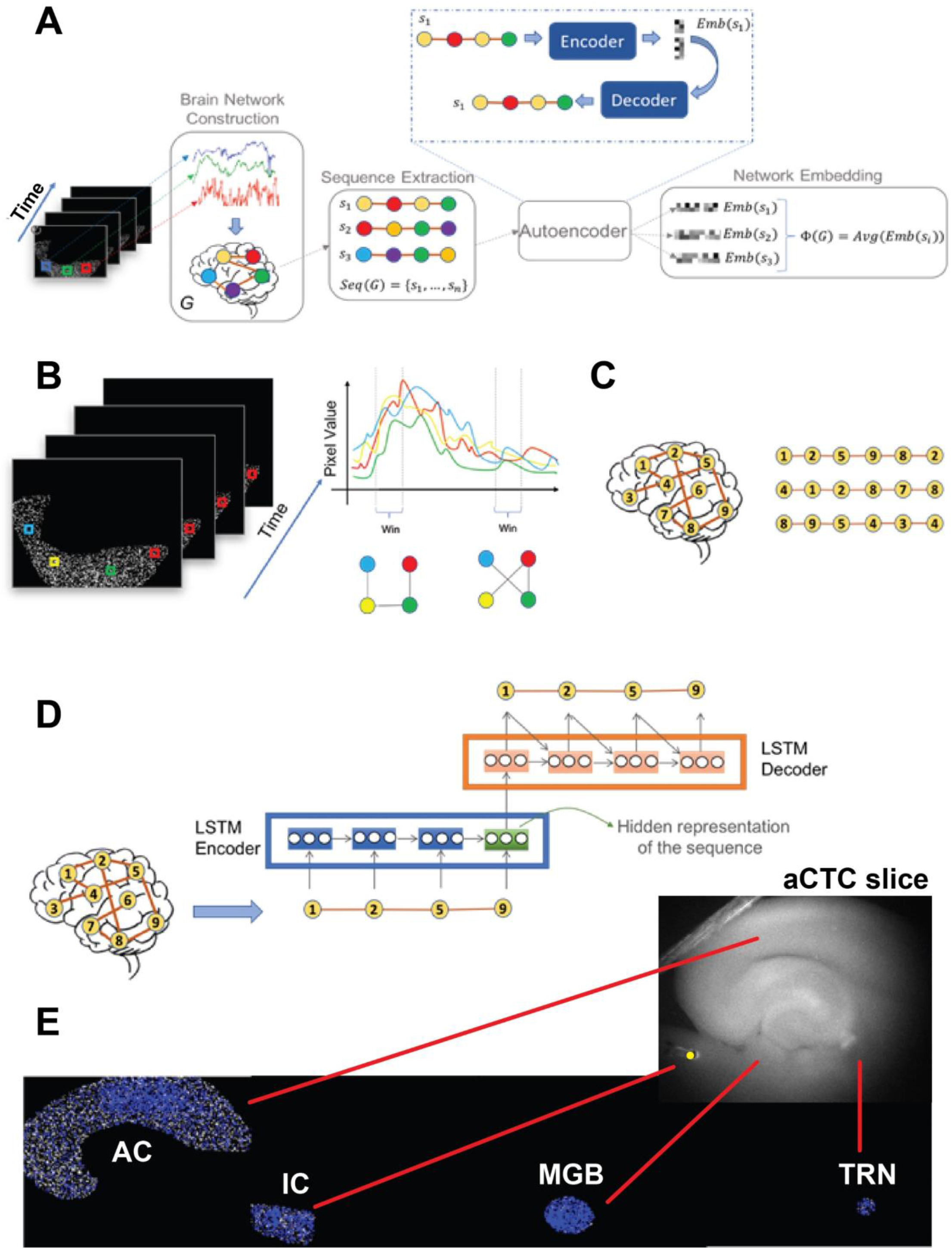
Procedures and output of brain network analysis: Cartoon images showing the procedures followed to run the brain network analysis on the FA imaging data obtained from AC, MGB, TRN, and IC of aCTC slice following the IC stimulation A) Proposed architecture for brain network representation learning, B) Time series shows the extraction of pixel values from the cortex images along the time axis followed by correlation network construction from time series. Each pixel is represented by a different color, C) Exemplary showing the sequence extraction from brain networks, D) The steps of sequence-to-sequence autoencoder, E) Examples of the correlation networks constructed from our data for activated areas of the brain structures, the blue points are the network’s nodes, Yellow circle: The site of the electric stimulation.

**Figure S3:**
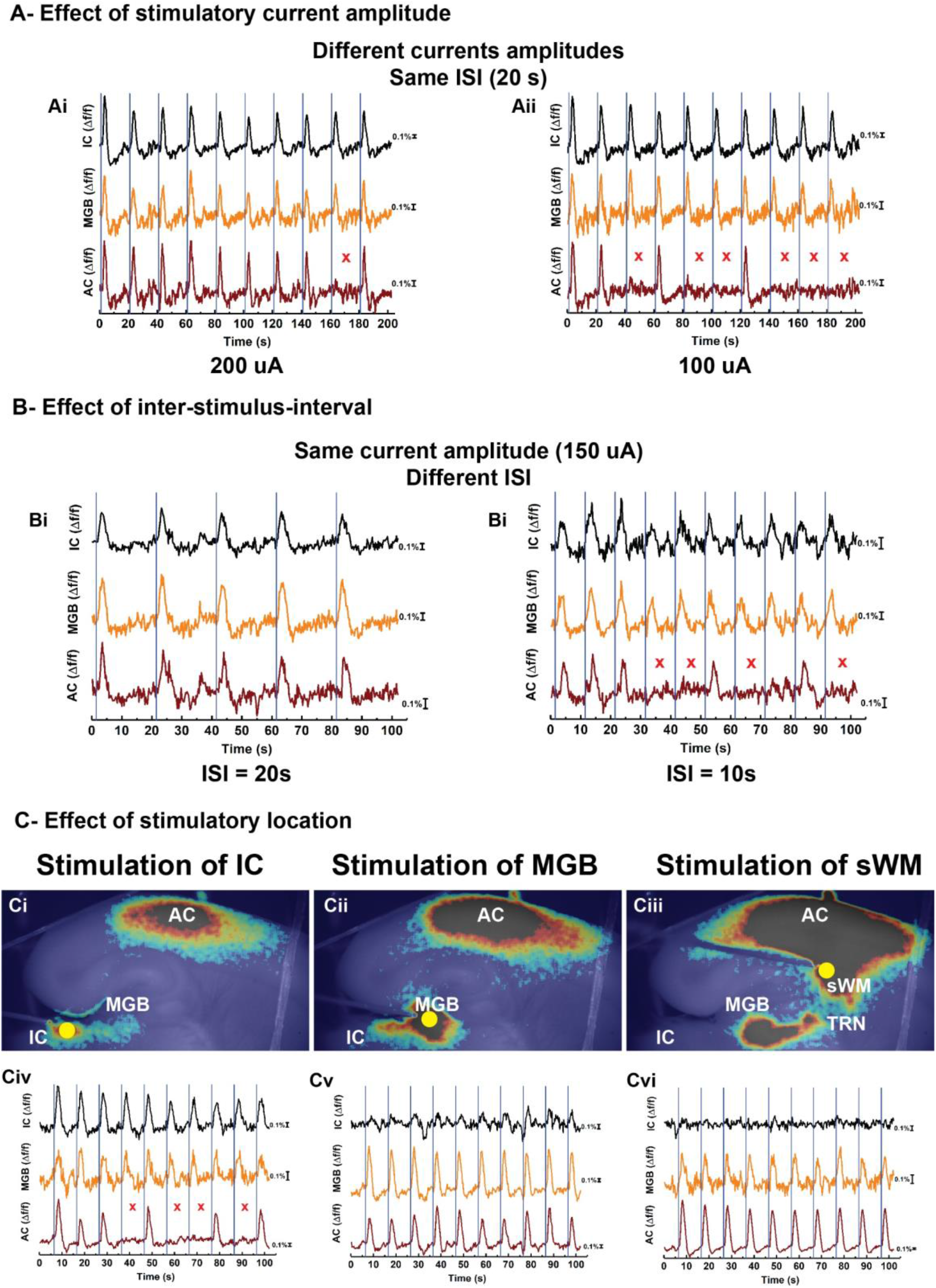
A) Ai and Aii: The time series of the Δf/f of the evoked FA signals of AC, MGB, and IC following stimulating the IC with 200 or 100 μA electric current, respectively, with the same ISI as 10 seconds, B) Bi and Bii: The time series of the Δf/f of the evoked FA signals of AC, MGB, and IC following stimulating the IC with 150 μA electric current with ISI = 20 or 10 seconds, respectively, C) Ci-Ciii: MATLAB pseudo-color images of the evoked FA signals in aCTC slice after the electric stimulation of IC, MGB, or the subcortical white matter, respectively, Civ-vi: The time series of Δf/f of the evoked FA signals in AC, MGB, and IC following the electric stimulation of IC, MGB, or subcortical white matter, red Xs refer to the occurrence of OFF cortical responses indicated by the absence of cortical FA, blue lines indicate the onset of the IC stimulation, yellow circles indicate the position of the electrical stimulation, ISI: Inter-stimulus interval, sWM: Subcortical white matter.

**Figure S4:**
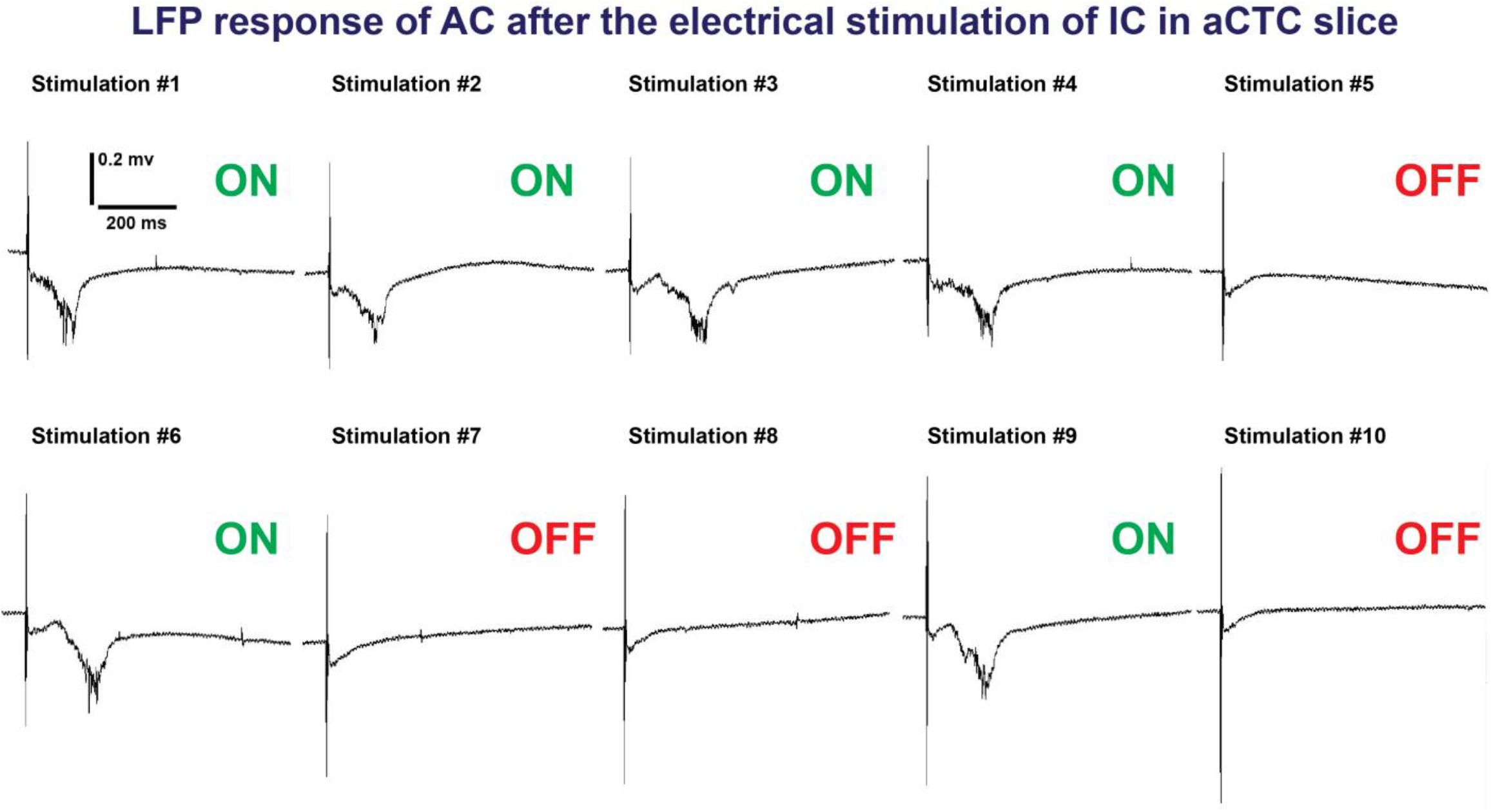
LFP signals recorded from cortical L3/4 after 10 trials of IC stimulation in a different laboratory environment (Matthew Banks laboratory, University of Wisconsin) using isoflurane anesthesia and no transcardiac perfusion, and still showing the binary cortical responses (ON vs OFF).

**Figure S5:**
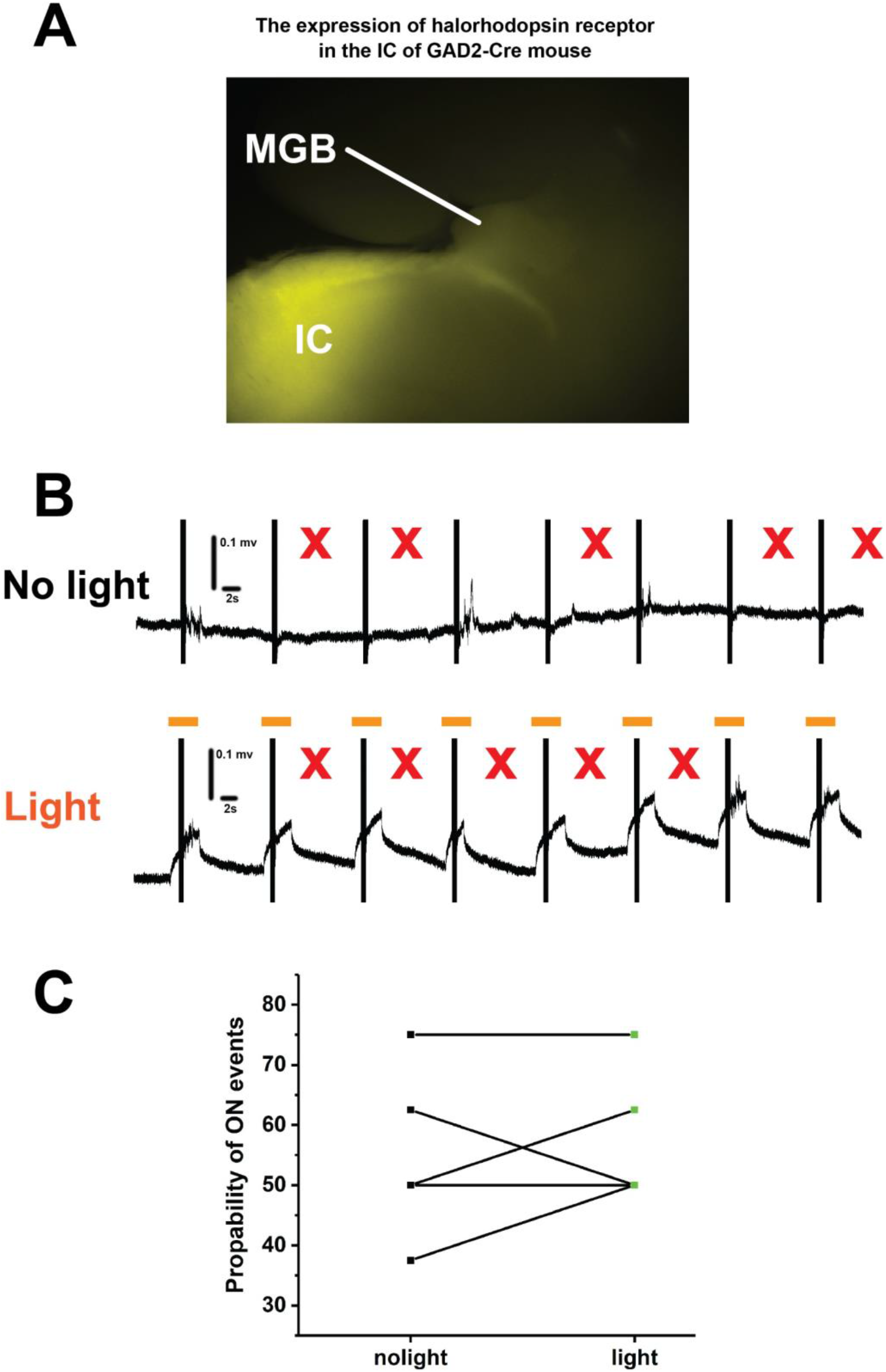
The OFF cortical responses were not driven by the feedforward inhibition of MGB by the GABAergic cells of the IC: A) Image of aCTC slice from GAD2-Cre mouse showing the Cre-dependent expression of halorhodopsin indicated by YFP tag in GABAergic cells of the IC as well as their projections to MGB, B) The time series of the post stimulus cortical LFP signals from L3/4 following the IC stimulation without (top) and with 565 nm light (bottom), C) A scatter plot of paired t-test showing no change in the probability of ON cortical responses after the photoinhibition of GABAergic cells by light, red Xs refer to the occurrence of OFF cortical responses indicated by the absence of post-stimulus cortical LFP signals from L3/4, orange lines indicate the time period of illuminating the light.

**Figure S6:**
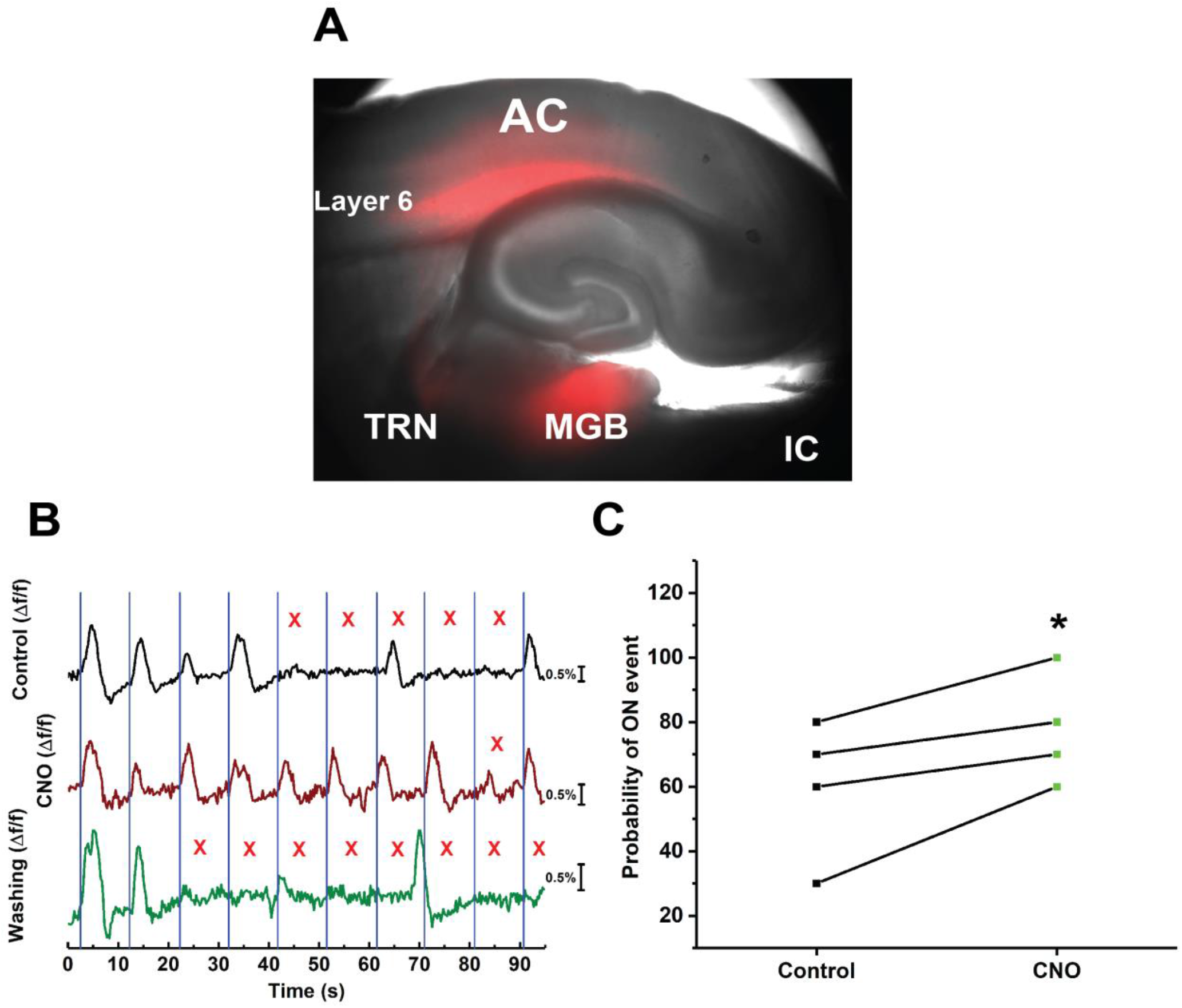
Chemical inhibition of CT-L6 cells retrieved the missing cortical responses. A) Image of aCTC slice of NTSR1-Cre mouse showing the expression of hM4Di receptors as indicated by m-cherry tag in NTSR1 +ve CTL6 cells as well as their projections to TRN and MGB, B) The time series of Δf/f of the evoked cortical FA signals during the perfusion of ACSF (black trace) and CNO (brown trace) then washing (green trace), C) A scatter plot graph of the paired t-test showing that the probability of ON cortical events was higher during the chemical inhibition of CTL6 cells by CNO compared to the control [*t(3)=-3.66, p=0.035], CNO: Clozapine-n-oxide.

**Figure S7:**
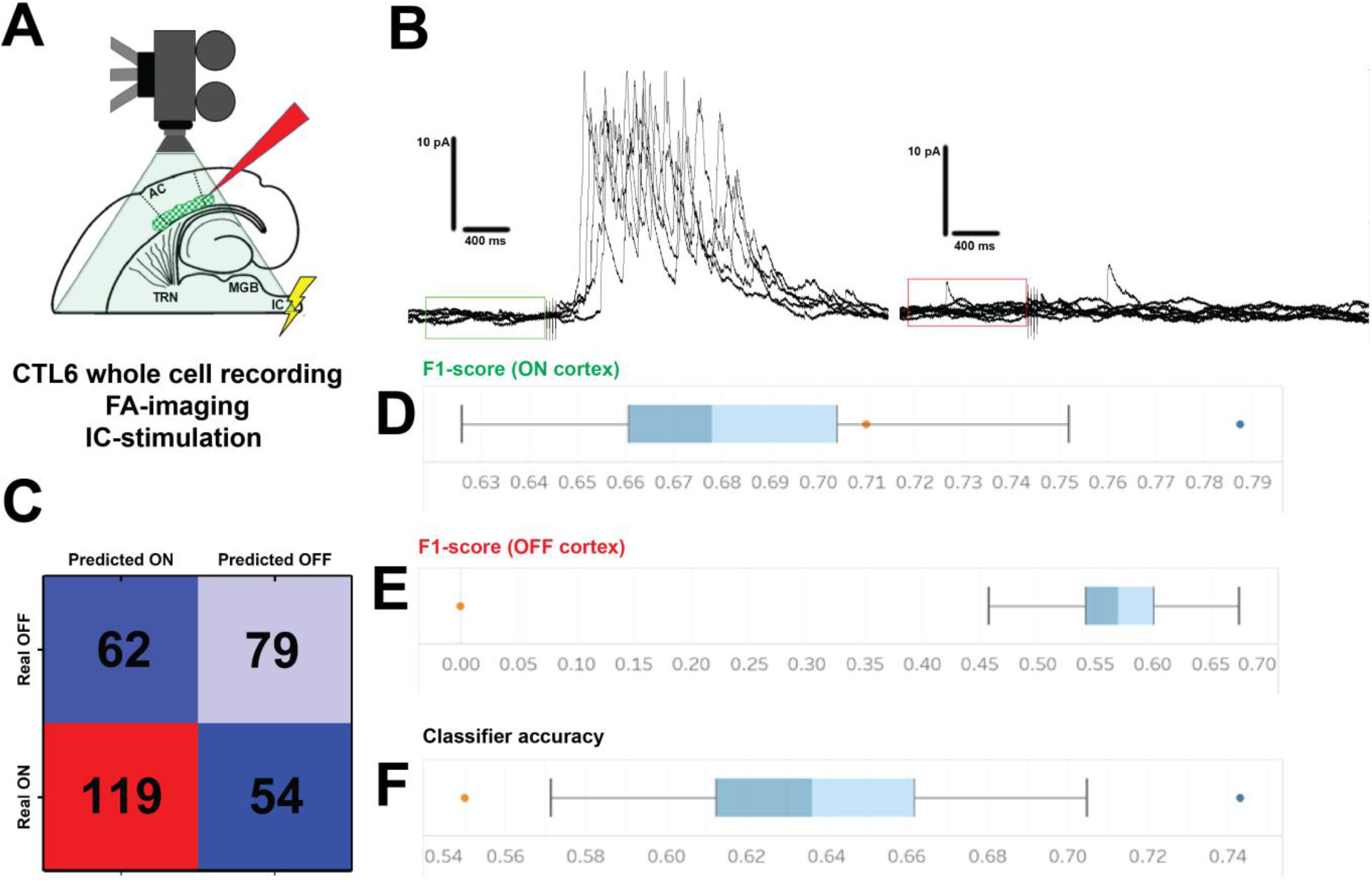
The oscillation of the pre-stimulus activity of CTL6 is a good predictor for the cortical response. A) A cartoon image showing the experimental design of simultaneous IC stimulation, FA imaging and whole cell recording from CT-L6 cells, B) Examples of multiple sweeps showing the pre-stimulus activity of CT-L6 cells during ON (left) vs OFF (right) cortical responses, green and red boxes assign the time period of one second before the stimulus onset that was taken for analysis under ON or OFF cortical responses, respectively, C) A confusion matrix showing the predicted ON vs OFF responses by the classifier against the real ON vs OFF, D) F1 score for the ON class of the classification compared to the majority classifier, represented with the orange point, E) F1 score for the OFF class of the classification compared to the majority classifier, represented with the orange point, F) Accuracy of the classification compared to the majority classifier, represented with the orange point, Blue points are the outliers of the result distribution, blue points are the outliers of the result distribution.

**Figure S8:**
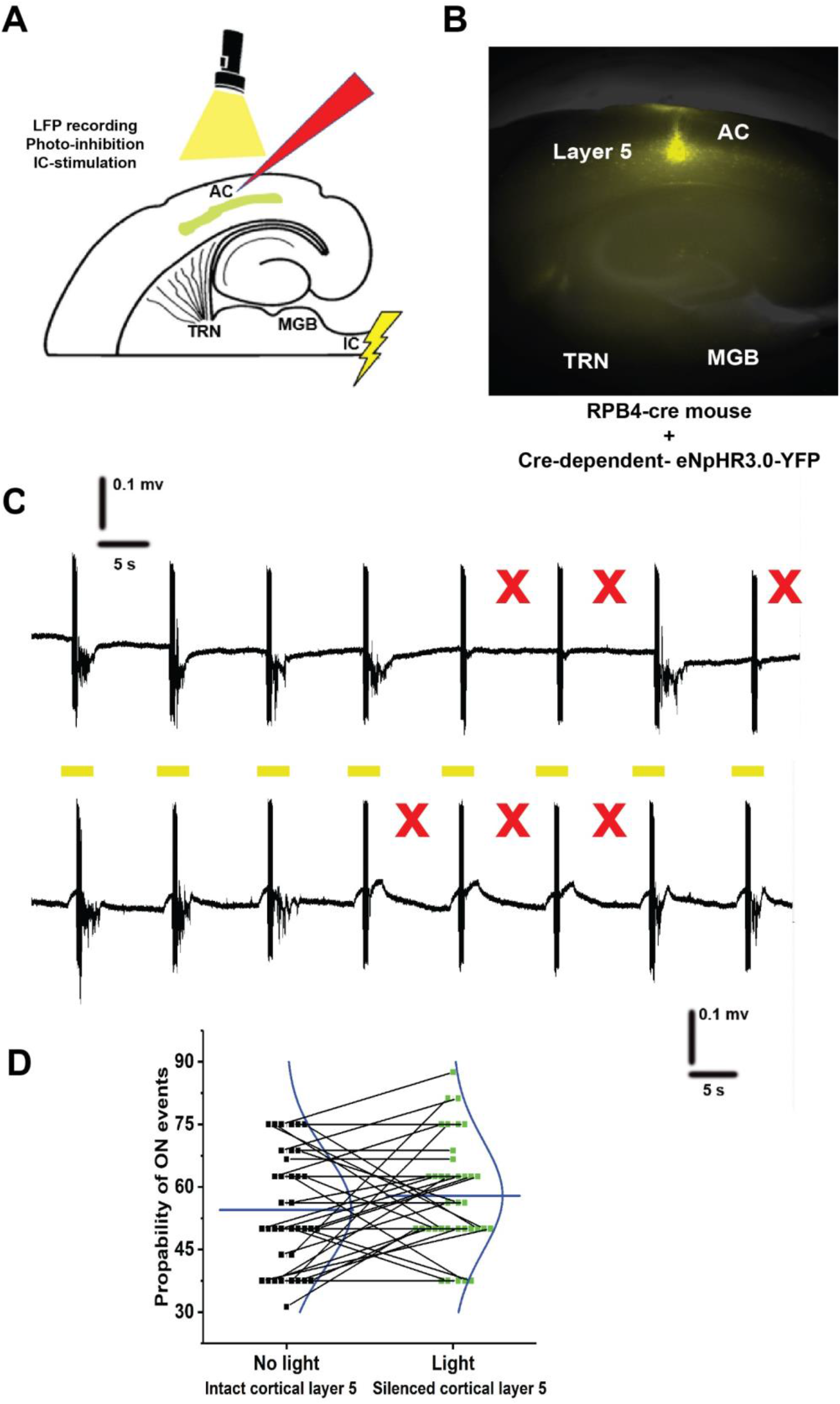
Photoinhibition of CT-L5 cells has no effect on the binary cortical response: A) A cartoon image showing the experimental design of simultaneous IC stimulation, LFP recording, and full field photoinhibition, B) Image of aCTC slice of RPB4-Cre mouse showing the expression of eNpHR3.0 receptors as indicated by YFP tag in RBP4 +ve L5 cells, C) The time series of the post stimulus cortical LFP signals from L3/4 following the IC stimulation without (top) and with 565 nm light (bottom), D) A scatter plot graph of paired t-test showing no difference in the probability of ON cortical responses during light or no light [paired t-test, t(37) = 0.46, p = 0.18], red Xs refer to the occurrence of OFF cortical responses indicated by the absence of the post-stimulus LFP signals recorded from L3/4, orange lines indicate the time period of light illumination.

